# Three new species, *Xanthomonas hawaiiensis* sp. nov., *Stenotrophomonas aracearum* sp. nov., and *Stenotrophomonas oahuensis* sp. nov., isolated from Araceae family

**DOI:** 10.1101/2023.09.17.558166

**Authors:** Shu-Cheng Chuang, Shefali Dobhal, Anne M. Alvarez, Mohammad Arif

**Author notes:** Repositories: The nucleotide sequences of 16S rRNA and *de novo* assembly genome sequences of four new species strains were deposited into the NCBI GenBank database under the following accession numbers. A5588^T^: OP964728 (16S rRNA) and CP115541-CP115542 (Bioproject: PRJNA914775) (whole genome sequence); A5586^T^: OP964727 (16S rRNA) and CP115543 (Bioproject: PRJNA914775) (whole genome sequence); A6251^T^: OP962219 (16S rRNA) and CP115873 (PRJNA916509); A2111: OP962220 (16S rRNA) and JAQMHB00000000 (PRJNA916509) (whole genome sequence).

## Abstract

*Xanthomonas* and *Stenotrophomonas* are closely related genera in the family Lysobacteraceae. In our previous study of aroid-associated bacterial strains, most strains isolated from anthurium, and other aroids were reclassified as *X. phaseoli* and other *Xanthomonas* species. However, two strains from *Spathiphyllum* and *Colocasia* were phylogenetically distant from other strains in the *Xanthomonas* clade and two anthurium strains clustered within the *Stenotrophomonas* clade. Phylogenetic trees based on 16S rRNA and nine housekeeping genes placed the former strains with type strain of *X. sacchari* from sugarcane and the latter strains with type strain of *S. bentonitica* from bentonite. In pairwise comparisons with type strains, the overall genomic relatedness indices required delineation of new species; digital DNA-DNA hybridization and average nucleotide identity values were lower than 70% and 95%, respectively. Hence, three new species are proposed: *S. aracearum* sp. nov. and *S. oahuensis* sp. nov. for two anthurium strains, and *X. hawaiiensis* sp. nov. for the spathiphyllum and colocasia strains. The genome size of *X. hawaiiensis* sp. nov. is ∼4.88 Mbp and higher than *S. aracearum* sp. nov. (4.33 Mbp) and *S. oahuensis* sp. nov. (4.68 Mbp). Pan and core-genome analyses revealed 426 and 576 core genes present in 40 xanthomonads and 25 stenotrophomonads, respectively. The average number of unique genes in *Stenotrophomonas* spp. was higher than in *Xanthomonas* spp. implying higher genetic diversity in *Stenotrophomonas*.

## Introduction

Genera *Xanthomonas* Dowson 1939 and *Stenotrophomonas* Palleroni and Bradbury 1993 are the groups of gram-negative and aerobic bacteria belonging to family Lysobacteraceae (syn. Xanthomonadaceae) of Lysobacterales (syn. Xanthomonadales) order of Gammaproteobacteria class in phylum Proteobacteria. *Stenotrophomonas* and *Xanthomonas* are phylogenetically closely related genera. Before the current generic name, *Xanthomonas* was described as *Bacterium* in 1921 and later reclassified into the genus *Phytomonas* in 1923 (Doidge 1921; Bergey et al. 1923; Dowson 1939). The taxonomy of the first reported xanthomonad from pepper and tomato was changed several times. The pathogen was originally classified as *B. vesicatorium* then *X. vesicatoria*; subsequently it was given the trinomial pathovar *vesicatoria* first under *X. campestris* (Young et al. 1978) and later under *X. axonopodis* (Vauterin et al. 1995), and finally reclassified as a separate species, *X. euvesicatoria* (Jones et al. 2004; Constantin et al. 2016). Interestingly, before designating the genus *Stenotrophomonas*, the first stenotrophomonad isolated from pleural fluid of a hospitalized patient had been referred to as *X. maltophilia*; initially identified within the *Bacterium* genus, it was subsequently reclassified under *Pseudomonas* for a decade (Hugh and Ryschenkow, 1961; Swings et al. 1983; Palleroni and Bradbury, 1993). In 1993, due to distinct phylogenetic lineage from the other phytopathogens in *Xanthomonas*, *X. maltophilia* was replaced by *S. maltophilia* (Palleroni and Bradbury, 1993). At the time of writing this manuscript, there are 36 and 17 validly published species of *Xanthomonas* and *Stenotrophomonas*, respectively, as well some other invalid species shown in quotation marks (“ “) throughout the rest of this article in the List of Prokaryotic names with Standing in Nomenclature (LPSN, last accessed Dec. 2022) (Parte et al. 2020).

Most of *Xanthomonas* species are pathogenic to more than 400 different monocot and dicot plants, including economically important crops and ornamentals. Additionally, some *Xanthomonas* strains are nonpathogenic and associated with plants (Bradbury, 1984; Leyns et al. 1984; Vauterin et al. 2000; Ryan et al. 2011; Vandroemme et al. 2013; Parte et al. 2020; Timilsina et al. 2020; Mafakheri et al. 2022). On monocotyledonous hosts, *X. oryzae* pv. *oryzae* and *X. oryzae* pv. *oryzicola* listed on the USDA Select Agent list and cause severe diseases of rice (*Oryza sativa*). Also, *X. albilineans* causes leaf scorch of sugarcane (*Saccharum officinarum*) and *X. vasicola* is a causal agent of banana wilt (Ryan et al. 2011). Whereas *X. maliensis*, “*X. sontii*”, “*X. indica*” were reported associated with the rice phytobiome (Triplett et al. 2015; Bansal et al. 2021; Rana et al. 2022), and *X. sacchari* was associated with sugarcane, and reported causing rice sheath rot disease (Ivayani et al. 2023). Constantin et al. (2016) reclassified bacterial strains isolated from *Anthurium*, *Dieffenbachia*, and other ornamental Araceae plants into the species and/or pathovars, *X. phaseoli* including two pathovars *dieffenbachiae* and *syngonii*, *X. citri* pv. *aracearum* and *X. euvesicatoria*. The Araceae strains of *X. phaseoli* pv. *dieffenbachiae*, *X. phaseoli* pv. *syngonii*, and *X. citri* pv. *aracearum* were pathogenic on their original hosts, but *X. euvesicatoria* strains isolated from *Philodendron* caused weak symptoms and lacked host specificity on tested Aracea hosts (Constantin et al. 2017).

As for *Stenotrophomonas* spp., they are ubiquitous environmental bacteria isolated from various sources. The type species, *S. maltophilia*, was an opportunistic pathogen on humans infecting through clinical materials and equipment, and “*S. sepilia*” was isolated from the nosocomial patient’s blood specimen (Hugh and Ryschenkow, 1961; Al-Anazi and Al-Jasser, 2014; Gautam et al. 2021). Besides the clinical species, *Stenotrophomonas* species are also from various sources, such as *S. humi* and *S. terrae* from soil, *S. daejeonensis* and *S. geniculata* from water, *S. pavanii*, *S. rhizophila*, and “*S. cyclobalanopsidis*” associated with plants, “*S. pennii*” and “*S. muris*” from animals, *S. lactitubi* and *S. indicatrix* from surfaces in contact with food, and *S. acidaminiphila* and *S. chelatiphaga* from sludges (Wolf et al. 2002; Heylen et al. 2007; Lee et al. 2011; Ramos et al. 2011; Bian et al. 2020; Gilroy et al. 2021; Afrizal et al. 2022; Weber et al. 2018; Assih et al. 2002; Kaparullina et al. 2009). Notably, *S. maltophilia* was also encountered frequently in aquatic and plant-associated environments, and *S. rhizophila* strains isolated from the rhizosphere and geocaulosphere were separated from *S. maltophilia* based on 16S rDNA analysis and DNA-DNA hybridization data (Denton and Kerr, 1998; Berg et al. 1996; Minkwitz and Berg 2001; Wolf et al. 2002).

The Araceae family, among monocot plants, includes the most economically important ornamental plants in Hawaii, especially the genus *Anthurium*. During the 1980s to 1990s, the anthurium industry was seriously damaged due to *X. phaseoli* pv. *dieffenbachiae* outbreak (formerly called *X. axonopodis* pv. *dieffenbachiae*) (Alvarez et al. 2006; Constantin et al. 2016). Hundreds of bacterial strains were isolated from various plant genera in Araceae worldwide including the strains collected during the outbreaks in Hawaii, and stored in the Pacific Bacterial Collection at University of Hawaii at Manoa (https://pacificbacterialcollection.com/). In our previous five-gene multilocus sequence analysis (MLSA) of Lysobacteraceae strains isolated from Araceae family, a strain from *Spathiphyllum* and another strain from *Colocasia* clustered within *Xanthomonas* clade but formed a distinct monophyletic lineage while two strains from *Anthurium* grouped with *Stenotrophomonas* clade instead of *Xanthomonas* clade (Chuang and Arif, unpublished). Moreover, these two stenotrophomonads were distinct from the other xanthomonads from Araceae based on the utilizations of N-acetyl-D-galactosamine (GalNAc) and D-serine, and the inabilities to oxidize D-galactose, glycerol, pectin, and sucrose based on Biolog GEN III microplate assays (Chuang and Arif, unpublished).

Hence, we sequenced the whole genomes of the former strains isolated from Araceae, which are potential novel species, and analyzed with the genomes of *Xanthomonas* spp. and *Stenotrophomonas* spp. type strains. Based on the nine-gene MLSA, overall genomic relatedness indices (OGRI) values, and pan-core genomic analyses, strains A6251^T^ from *Spathiphyllum* and A2111 from *Colocasia* are described as new species *X. hawaiiensis* sp. nov., strain A5588^T^ from *Anthurium* as *S. aracearum* sp. nov., and strain A5586^T^ from *Anthurium* as *S. oahuensis* sp. nov.

## Methods

### Bacterial DNA Isolation and Genome Sequencing

Bacteria were streaked out from the culture stock and grew on 2, 3, 5-triphenyltetrazolium chloride (TZC) agar medium (dextrose 5 gL^-1^, peptone 10 gL^-1^, 0.001% sterilized TZC, and agar 18 gL^-1^) at 28 °C for 2 days. Bacterial genomic DNA was isolated from pure culture using QIAGEN Genomic-tip 100/G following the manufacturer’s instruction of (Qiagen, Valencia, CA). The Seqwell plexWell LP384 Library Preparation Kit and Native Barcoding Kit 24 V14 (SQK-NBD112.24) were used for barcode-indexed whole genome sequencing with Illumina NovaSeq system (Illumina San Diego, USA) and Oxford Nanopore MinIoN Mk1C device (Oxford Nanopore Technologies, ONT, Oxford, UK), respectively. ONT long reads were base called and demultideplexed using basecaller and barcoder of GUPPY v6.3.2 on MANA, a high-performance computing cluster at the University of Hawaii at Manoa.

### Hybrid Genome Assembly and Genome annotation

The hybrid assembler, Unicycler v0.4.8 plugged in the web-server of BV-BRC 3.26.4 (https://www.bv-brc.org/), was employed by uploading paired-end (2 x 150bp) Illumina short reads and high-accuracy basecalled ONT long reads for *de novo* genome assemblies (Wick et al., 2017; Wattam et al., 2017; Klair et al, 2022; Olson et al., 2023). Briefly, Unicycler carried out SPAdes (v3.13.0) to assemble the Illumina short reads and then ran miniasm, minimap2 (v2.17), and Racon (v1.4.13) for long-read plus contig assembly, long-read bridging, and contig polishing. Alternatively, the genome of the strain (A2111) with a lower coverage of short reads was assembled by performing Flye v2.9.1 and genome assembly pipeline in the web server of BV-BRC 3.26.4 (https://www.bv-brc.org/). The genome annotations were performed using Prokaryotic Genome Annotation Pipeline (PGAP v4.10) on NCBI (Tatusova et al. 2016) and Rapid Annotation using Subsystem Technology (RAST v2.0) web server (Aziz et al. 2008) as well.

### Phylogenetic Analyses

The partial 16S rRNA gene sequences of new species strains were amplified using primer set P16S-F1 (5’-AGACTCCTACGGGAGGCAGCA-3’) and P16S-R1 (5’-TTG ACGTCATCCCCACCTTCC-3’) by end-point PCR (Larrea-Sarmiento et al. 2019). Each 25 µl of PCR reaction mix contained 5 µl of 5X Q5 buffer, 5 µl of GC enhance, 2.5 µl of 5 µM primer F and R, 0.5 µl of 2.5 mM dNTPs, 0.5 µl of Q5 polymerase, 1 µl of gDNA, and 6.5 µl of nuclease-free water. The PCR reaction was run following program: 10 sec at 98°C; 35 cycles of 10 sec denaturing at 98°C, 30 sec annealing at 58°C, and 30 sec extending at 72°C; 2 min at 72°C for final extension in a T100 Thermal Cycler (BIO-RAD Lab. Inc., Hercules, CA). The sizes of PCR products were checked by running agarose gel electrophoresis, purified using Exo (exonuclease I) –SAP (shrimp alkaline phosphatase) (GE Healthcare, Little Chalfont, UK) method by following the manufacturer’s instruction, and sent for Sanger sequencing service at the GENEWIZ company (South Plainfield, NJ). The sequences were double checked with the assembled genomes.

For the phylogenetic analysis of 16S rRNA gene, the full-length sequences of 65 *Stenotrophomonas* and *Xanthomonas* species type strains were retrieved from their whole genome sequences downloaded from the GenBank on NCBI (Table S1). The multiple aligned was performed using Geneious Prime 2021.2.2 (http://www.geneious.com). The module of finding the best DNA/Protein model for the multiple alignment data was conducted, and the Maximum Likelihood (ML) phylogenetic tree was built using MEGA X (Kumar et al. 2018). The consistency of the phylogenetic tree was assessed by computing 1,000 bootstrapping analyses.

Additionally, the precise MLSA was performed to reveal the phylogenetic relations among the novel species and other *Stenotrophomonas* and *Xanthomonas* species. Nine housekeeping genes (*atpD*, *dnaA*, *dnaK*, *gltA*, *gyrB*, *nuoD*, *ppsA*, *rpoH*, and *uvrB*), used from the previous studies (Ramos et al. 2011; Vasileuskaya-Schulz et al. 2011; Chuang and Arif, unpublished) were retrieved from downloaded genomes. The sequences of the nine housekeeping genes were aligned with free end gaps algorithm, trimmed both ends, and concatenated in alphabetic order for analyzing using Geneious Prime. The ML phylogenetic tree was formed using MEGA X following the process as detailed above. The phylogenetic trees with bootstrapping analyses were created using web-based tool Interactive Tree Of Life (iTOL v6, https://itol.embl.de).

### Genome Similarity

In order to define new species, the pairwise comparisons of overall genomic relatedness indices (OGRIs) among the genomes of new species strains and other type strains of *Stenotrophomonas* and *Xanthomonas* species retrieved from NCBI database were calculated. The pairwise ANI and AP (alignment percentage) values were calculated using CLC Genomics Workbench 22.0.2 (CLC Bio-Qiagen, Arahus, Denmark). Due to inclusion of some incomplete genomes, OrthoANI (Average Nucleotide Identity by Orthology), which only considered the orthologous fragment pairs, was additionally calculated by performing Orthologous Average Nucleotide Identity tool (OAT) (Lee et al. 2016). Moreover, the pairwise dDDH values and the differences in G+C content (mol%) were inferred by estimating precise distance from whole genome sequences using the Genome-Genome Distance Calculator (GGDC) v3.0 on Type Strain Genome Server (TYGS) web server (https://tygs.dsmz.de/) (Meier-Kolthoff et al. 2013; 2022).

### Pan-Genome Analysis

Whole genome sequences of the new species and closely phylogenetically related species in each genus were used for pan– and core-genome analyses. The Prokka v1.14.6 (Seemann, 2014) was used to re-annotate representative genomes, and the output gff files were used as input files for the Roary v3.13.0 pipeline (Page et al. 2015). For Roary, core and accessory genes were assessed with 80% minimum BLASTp identity and multi-FASTA alignment of the core genome was generated using highly accurate PRANK, which is a probabilistic multiple alignment program (Löytynoja, 2014; Page et al. 2015). The number of core and unique genes among species of each genus were assessed from the Roary output and were used for the flower plots by computing R script in RStudio. A core gene phylogenetic tree was established using a ML tree inference tool Randomized Axelerated Maximum Likelihood – Next Generation (RAxML–NG) v0.8.0 (Kozlov et al. 2019), which combine the strengths of RAxML (Stamatakis, 2014) and Exascale Maximum Likelihood (ExaML) (Kozlov et al. 2015). The DNA substitution model, General Time Reversible (GTR) + GAMMA (G), was performed and ran separately with core genomes of type species of *Xanthomonas* spp. and *Stenotrophomonas* spp. with 1,000 bootstrap replicates. The core genome phylogenetic tree was displayed using web-based tool Interactive Tree Of Life (iTOL v6, https://itol.embl.de). The Roary matrix with the present and absence of core and accessory genes was combined with core genome ML tree and the results were visualized by conducting roary_plots.py (Page et al. 2015).

### Antibiotic Sensitivity Assay

Antibiotic sensitivity assays were performed using disc diffusion method described in Klair et al. (2022). Single colony was picked from the pure culture plates of four new species strings and incubated in 10 ml of LB broth and incubated at 28°C with 200 rpm shaking for 16 hours. Light absorbance at 600nm (OD600) of bacterial inoculum was adjusted to the value about 1.0 and 100 µl of inoculum was spread evenly on Nutrient agar (NA, CRITERION^TM^, Hardy Diagnostics). Seven antibiotics with different concentrations including Bacitracin (50 mg/ml), Chloramphenicol (50 mg/ml), Gentamicin (50 mg/ml), Kanamycin (50 mg/ml), Penicillin (50 mg/ml), Tetracycline (40 mg/ml), and Polymyxin B Sulfate (50 mg/ml) were tested. One petri dish was divided into four zones and three discs impregnated with each antibiotic solution and one disc soaked with sterile distilled water as control were placed in the center of each zone. Inhibition zones were observed and measured the size after incubating the plates at 28°C for 24 hours.

## Results

### Genome Assembly and Annotation

The high-quality genome of the strains A6251^T^, A5588^T^ and A5586^T^ were assembled using Unicycler v0.4.8, whereas the strain A2111 had a better *de novo* assembly using another hybrid genome assembler, Flyev2.9.1 (Table 1). The genome sizes of new species strains from anthurium, A5588^T^ and A5586^T^, are of 4.33 Mbp and 4.68 Mbp with 66.44 mol% and 65.3 mol% of GC content, respectively. Meanwhile, the GC content were higher, in other two strains, *i.e.* A6251^T^ (4.88 Mbp) from spathiphyllum and A2111 (4.87 Mbp) from colocasia, with 68.93 and 68.88 mol% GC content, respectively (Table 1). Based on the annotation of NCBI-PGAPservice, the average CDS number of four strains was 4,016. The strain A5588^T^ has the least CDS number, whereas the strain A5586^T^ possessed the most number (Table 1). Although the CDS numbers estimated by RAST web server were slightly different from PGAP annotation, strain A5588^T^ had the lowest CDS number correlated with its genome size (data not shown). By contrast, the coverage of subsystem features presented in A5588^T^ was the highest and in A5586^T^ was the lowest (Figure 1A). Four strains comprised 23 out of the total 27 subsystem feature categories including virulence, stress response, membrane transport, DNA and protein metabolism (Figure 1B). Notably, only strains A6251^T^ and A2111 contained proteins in the iron acquisition and metabolism subsystem but not strains A5588^T^ and A5586^T^ (Figure 1B). *Stenotrophomonas maltophilia* was reported to use two putative iron acquisition systems for the mediation of siderophore and heme as iron starvation (Kalidasan et al. 2018), implying that strains A5588^T^ and A5586^T^ from anthurium were different from opportunistic human pathogen *S. maltophilia*.

**Figure 1.**
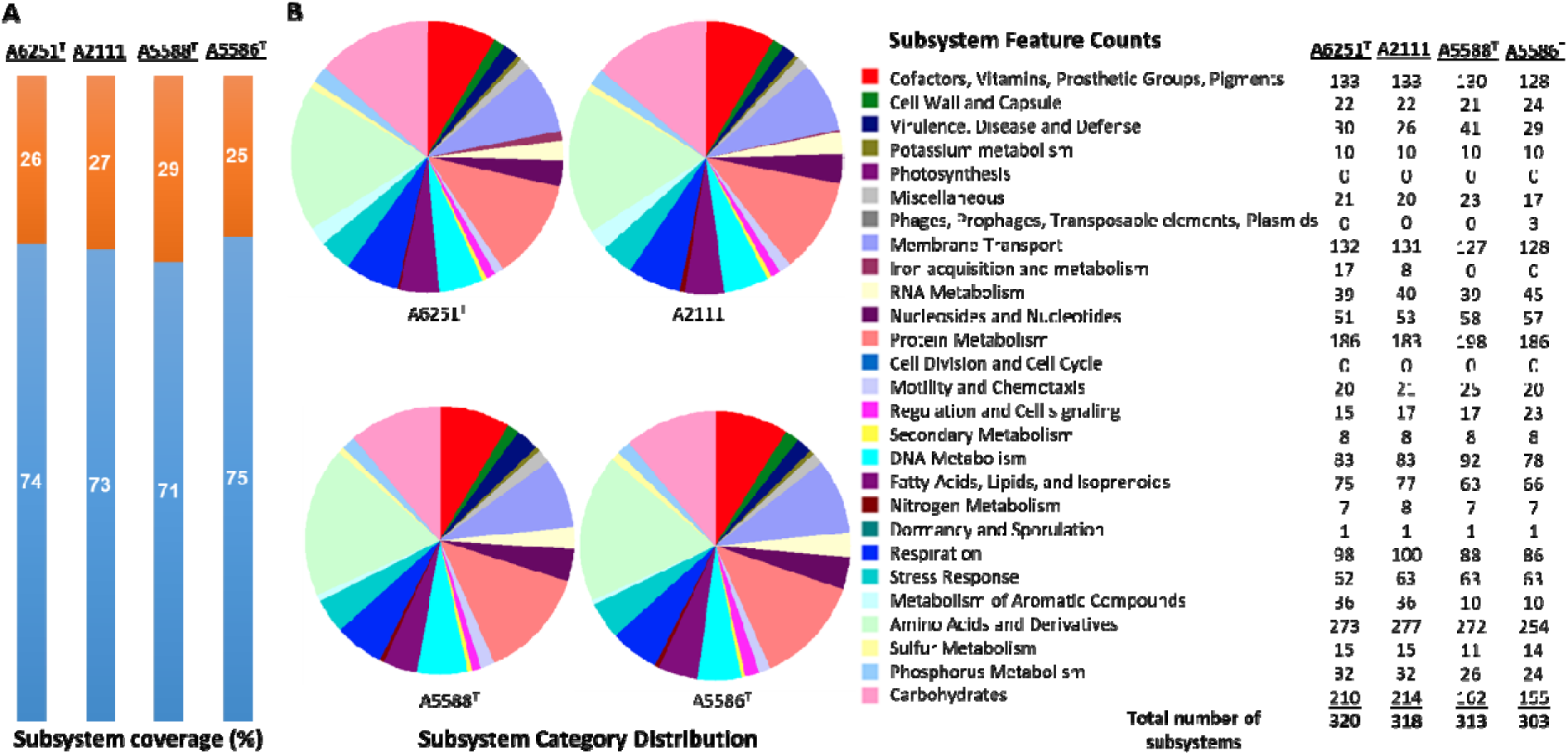
Subsystem annotation summary of new species strains, A6251^T^, A2111, A5588^T^, and A5586^T^ by conducting RAST web server. (A) The percentages of protein-coding genes present (orange portions) or absent (blue portions) in the RAST subsystem. (B) The pie chart and the number of subsystem features in total 27 categories found in four genomes of aroid strains.

**TABLE 1.** Genome features of *Xanthomonas hawaiiensis* sp. nov., *Stenotrophomonas aracearum* sp. nov., and *S. oahuensis* sp. nov. in this study.

| Species Name | Genome Size (Mbp) | No. of Contigs | N50 (bp) | GC content (%) | No. of CDS* | Short read average coverage | Long read average coverage | Genome Assembler | NCBI Accession no. |
| --- | --- | --- | --- | --- | --- | --- | --- | --- | --- |
| *Xanthomonas hawaiiensis* A6251<sup>T</sup> | 4.880993 | 1 | 4,880,993 | 68.93 | 4087 | 203.00 | 20.75 | Unicycler v0.4.8 | CP115873 |
| *Xanthomonas hawaiiensis* A2111 | 4.867870 | 3 | 4,828,414 | 68.88 | 4085 | 184.95 | 21.43 | Flye v2.9.1 | JAQMHB000000000 |
| *Stenotrophomonas aracearum* A5588<sup>T</sup> | 4.328236 | 1 | 4,328,236 | 66.44 | 3769 | 287.19 | 53.82 | Unicycler v0.4.8 | CP115543 |
| *Stenotrophomonas oahuensis* A5586<sup>T</sup> | 4.684619 | 2 | 4,623,839 | 65.30 | 4121 | 243.04 | 43.90 | Unicycler v0.4.8 | CP115541-CP115542 |

\* indicates the numbers of estimated CDS were generated from the NCBI PGAP pipeline.

### Phylogenetic Analyses

The partial sequences of 16S rRNA gene were amplified using primer set, P16S-F1 and P16S-R1, and deposited in the NCBI GenBank database under the accession number OP962219 (A6251^T^), OP962220 (A2111), OP964727 (A5586^T^), and OP964728 (A5588^T^). The 16S rRNA gene sequences were retrieved from the whole genomes of new species strains, and 38 type strains of *Xanthomonas* species and 23 type strains of *Stenotrophomonas* species published in NCBI database (Table S1). The sequences of the nearly entire 16S rRNA gene ranging from 1,415 bp (*S. bentonitica* LMG 29893^T^) to 1,421 bp (*S. chelatiphaga* DSM 21508^T^) were analyzed for phylogenetic relationships. The 16S rRNA gene sequences of A6251^T^and A2111 were identical with *X. sacchari* CFBP4641^T^ and only one base difference from “*X. sontii*“ PPL1^T^. The A5588^T^ and A5586^T^ were closely related to each other and *S. bentonitica* LMG 29893^T^ and showed higher similarity values ranging from 99.6 to 99.8% (Table 3). In the maximum likelihood (ML) phylogenetic tree, 16S rRNA gene sequences depicted better resolution within *Stenotrophomonas* species than *Xanthomonas* species because of very poor species discrimination, which was higher than 98.7% cutoff of 16S similarity (Figure 2).

**Figure 2.**
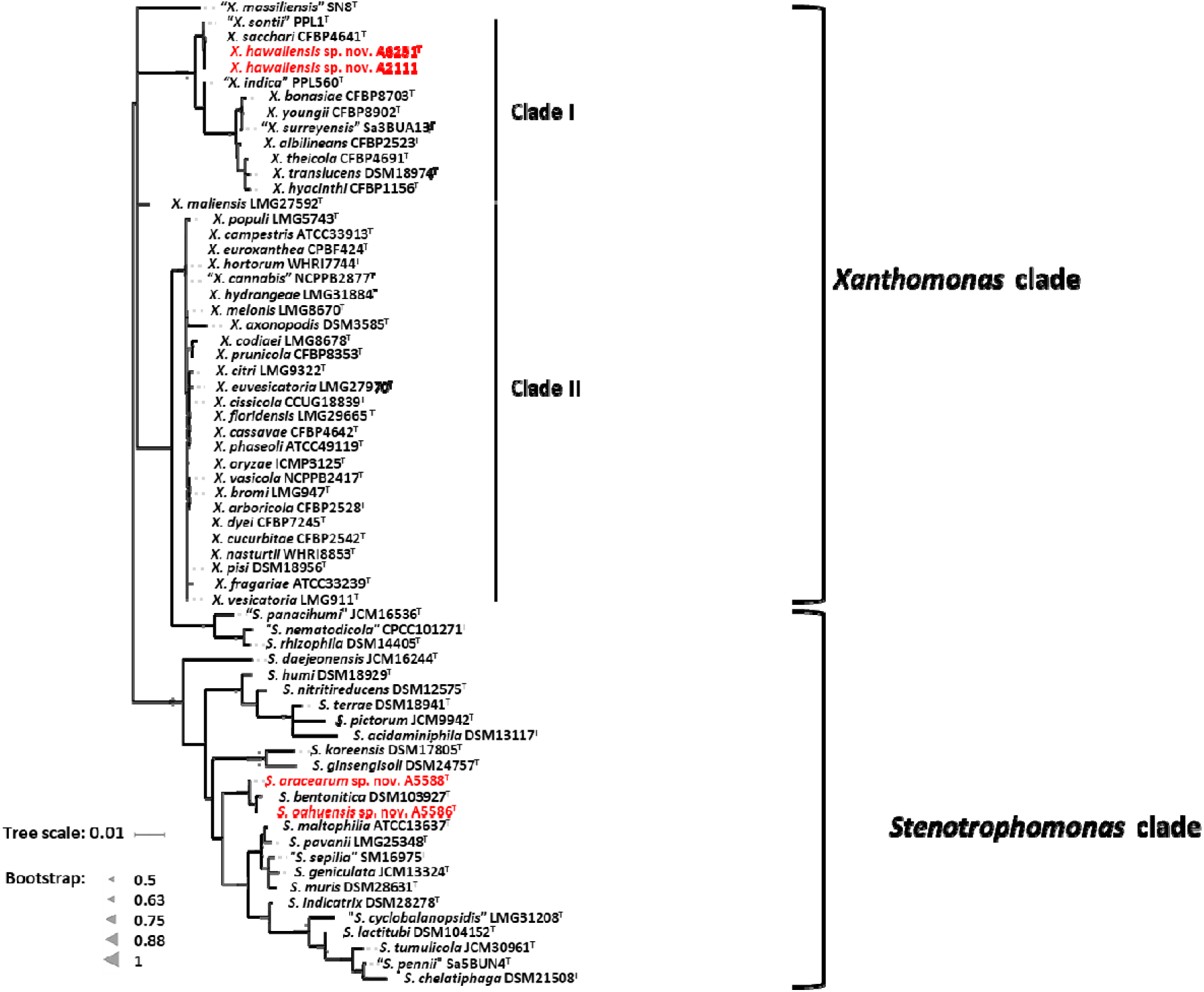
Maximum Likelihood phylogenetic tree based on almost full-length 16S rRNA gene sequences among three new species strains and type strains of *Xanthomonas* and *Stenotrophomonas* species. The tree scale bar indicates the number of nucleotide substitutions per sequence position. The range of gray triangles represent the degree of bootstrapping values.

For more detailed phylogenetic analysis, nine housekeeping genes (*atpD*, *dnaA*, *dnaK*, *gltA*, *gyrB*, *nuoD*, *ppsA*, *rpoH*, and *uvrB*) were selected and retrieved from whole genomes of formerly mentioned type strains of *Xanthomonas* and *Stenotrophomonas* species. Total length of concatenated sequence with nine genes in alphabetic order was about 14.3 Kb, which contained the maximum ∼2,443 bp of *gyrB* gene and the minimum ∼879 bp of *uvrB* gene sequences. The similarity of the concatenated gene sequences of two strains A6251^T^ and A2111 was 99.3%; strain A5588^T^ and strain A5586^T^ showed 89.8% similarity. The ML tree based on nine housekeeping genes indicated that two major phylogenetic clades, Clade I and Clade II, were present within the *Xanthomonas* clade with high bootstrapping value support (Figure 3). Similar Clade I and II phylogenetic groupings were reported in the previous studies (Koebnik et al. 2021; Mafakheri et al. 2022; Rana et al. 2022). The strains A6251^T^ and A2111 formed a monoclade clustering with *X. sacchari*, *X. indica*, *X. sontii*, *X. albilineans* in the Clade I, which also includes *X. surreyensis*, *X. bonasiae*, *X. traslucens*, *X. hyacinthi*, *X. theicola*, and *X. youngii* (Figure 3). *Stenotrophomonas bentonitica* consistently clustered with strains A5588^T^ and A5586^T^ with strong bootstrapping value. *Stenotrophomonas rhizophila* and *S. nematodicola* formed a clade closely related to the A5588^T^-A5586^T^-*S. bentonitica* clade, however, no grouping was formed in either tree (Figure 2 and 3).

**Figure 3.**
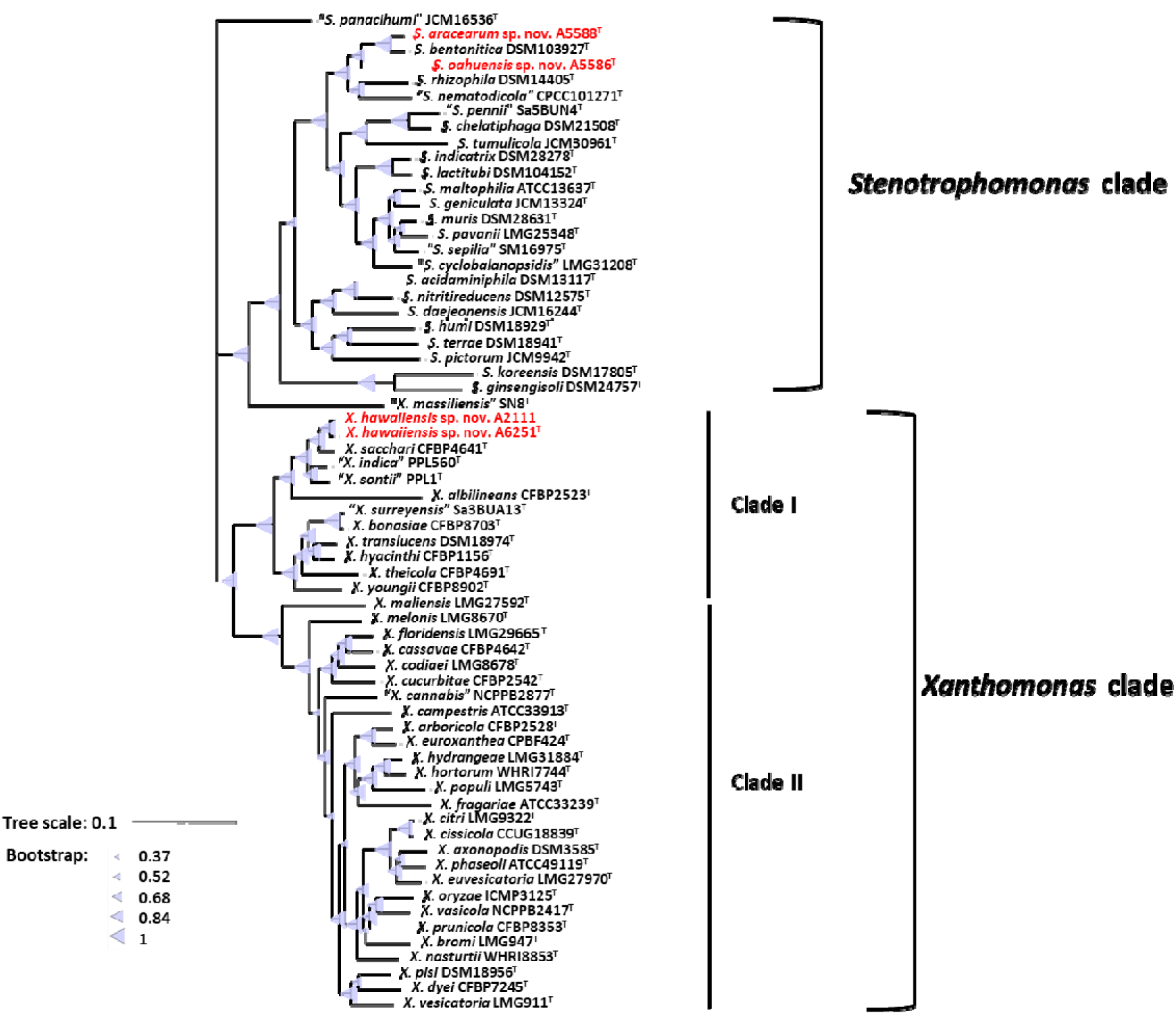
Maximum Likelihood phylogenetic tree based on concatenated sequence set of nine housekeeping genes, *atpD*, *dnaA*, *dnaK*, *gltA*, *gyrB*, *nuoD*, *ppsA*, *rpoH*, and *uvrB* of *Xanthomonas* and *Stenotrophomonas* species type strains. The scale bar represents the nucleotide substitutions per site. The range of purple triangles indicate the degree of bootstrapping support.

### Overall Genomic Relatedness Indices

In order to examine the accurate taxonomic classification, the overall genomic relatedness indices (OGRIs) including the values of ANI and dDDH of A6251^T^ from spathiphyllum and A2111 from colocasia were analyzed with other type strains of *Xanthomonas* spp., meanwhile A5588^T^ and A5586^T^ strains were compared with other type strains in *Stenotrophomonas*. The general cutoff values of ANI and dDDH for species delineation are lower than 95-96% and 70%, respectively (Goris et al. 2007; Richter 2009; Meier-Kolthoff et al. 2013).

Strains A6251^T^ and A2111 shared 98.4% ANI and 85.2% dDDH with each other, which indicated that two strains belong to the same species. Based on the pairwise comparisons of the other *Xanthomonas* spp. reference genomes with either A6251^T^ or A2111, the ANI and dDDH values were 83.4-94.9% and 22.3-59.3%, respectively, strongly signified that A6251^T^ and A2111 are distinguished from the others and should be considered a novel lineage (Table 2). Despite that *X. sacchari* CFBP4641^T^ shared slightly higher OrthoANI values (95.04, 95.1) with A6251^T^ and A2111, other OGRIs supported the assignment as a new species (Table 2).

**TABLE 2.**
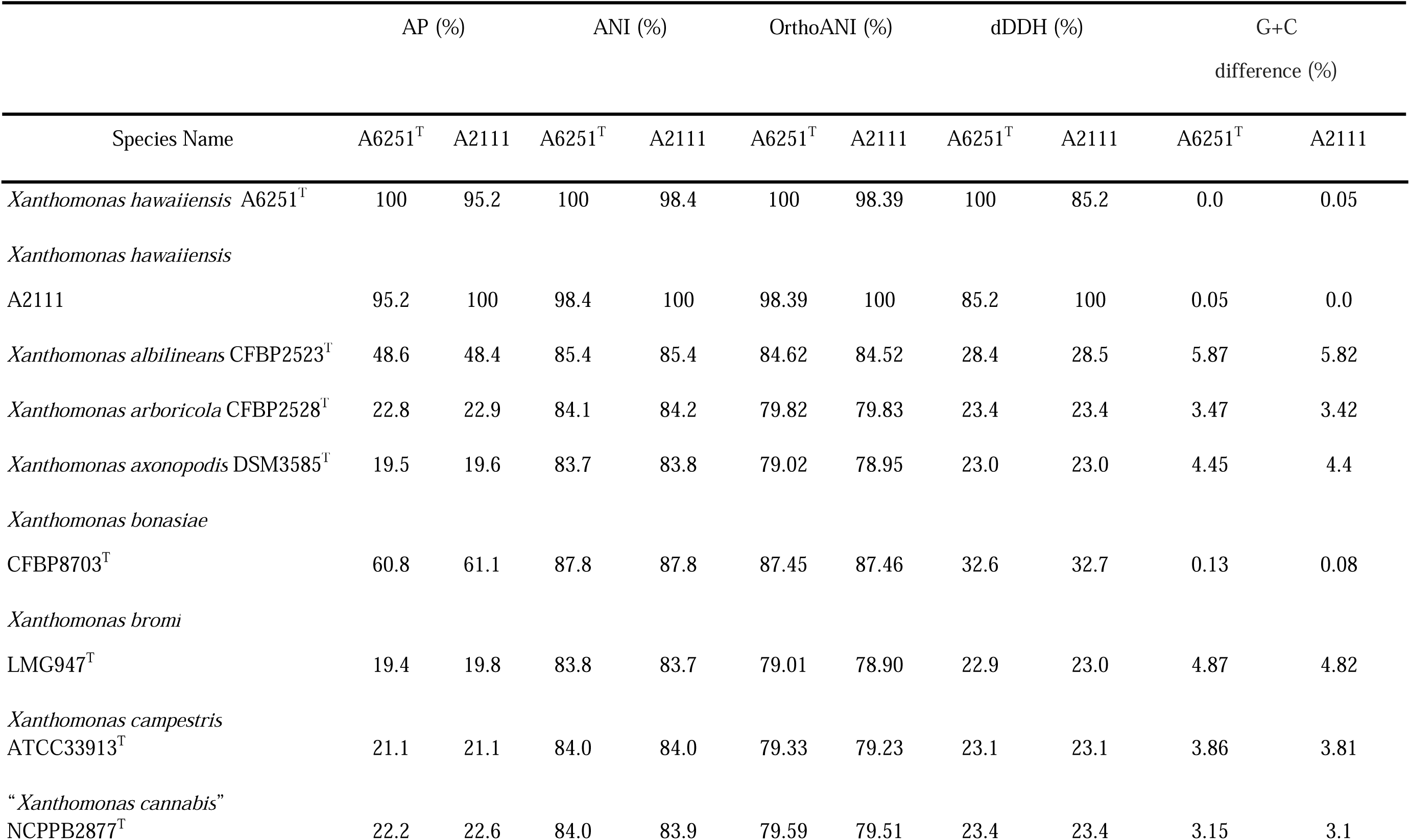
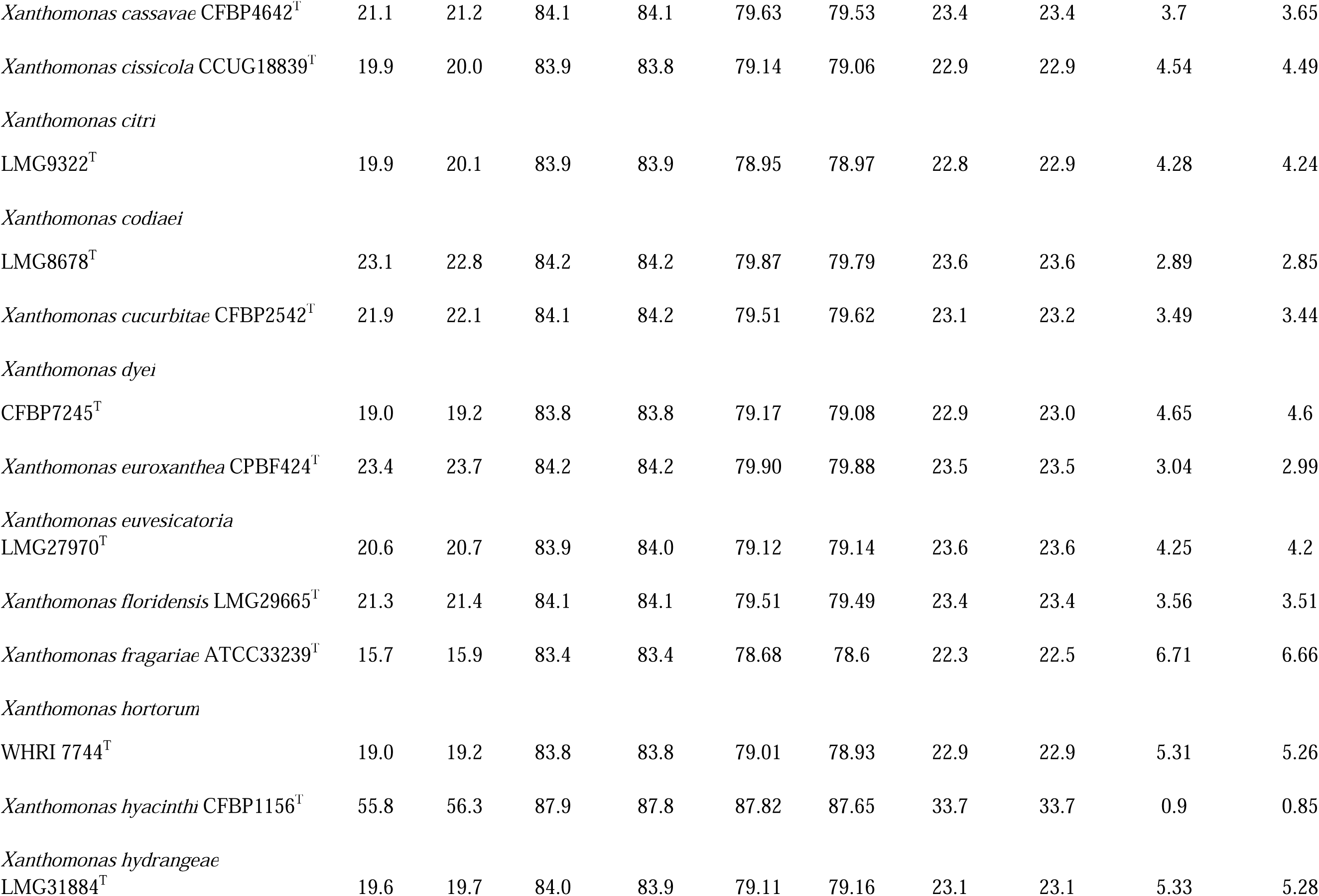
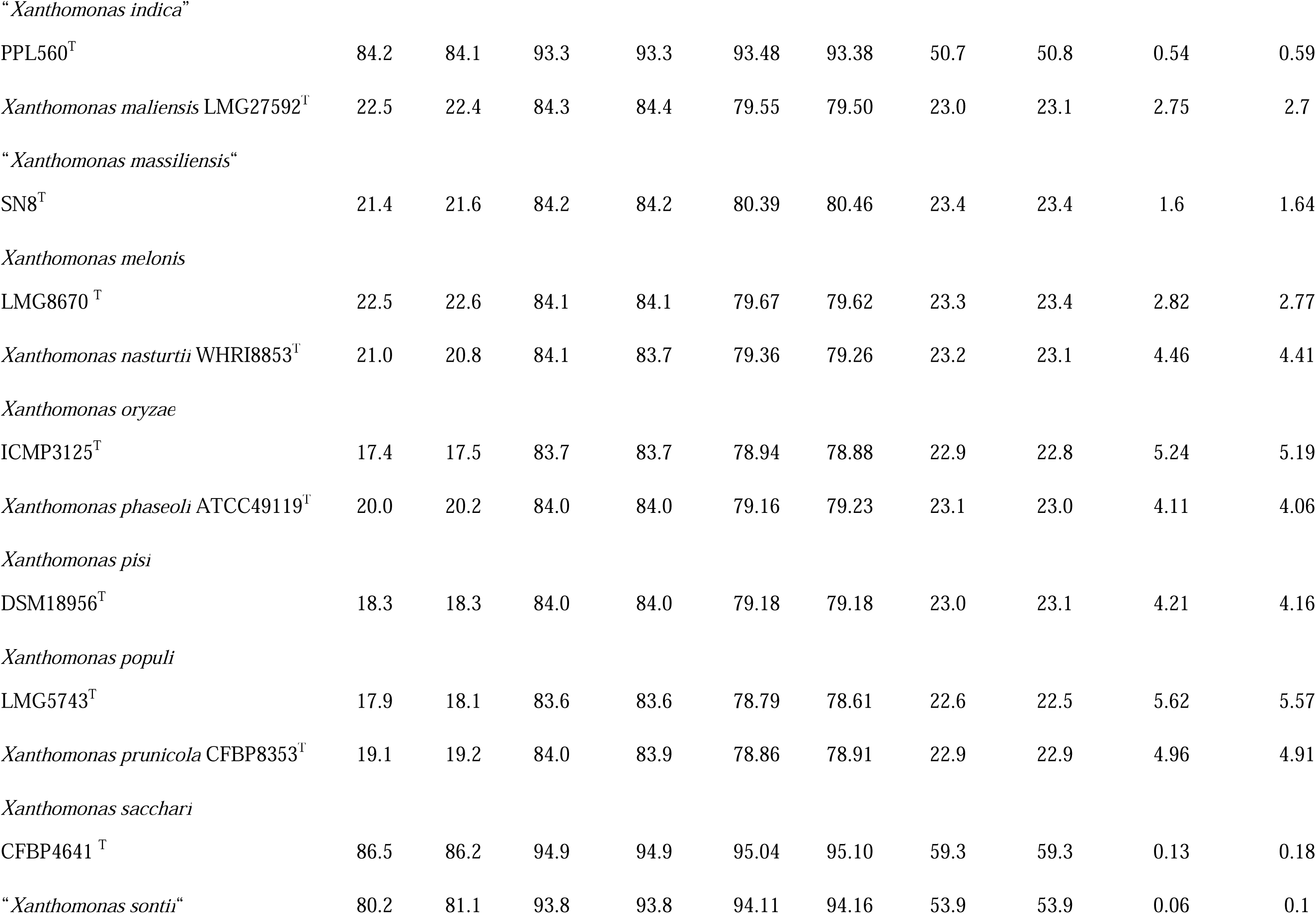
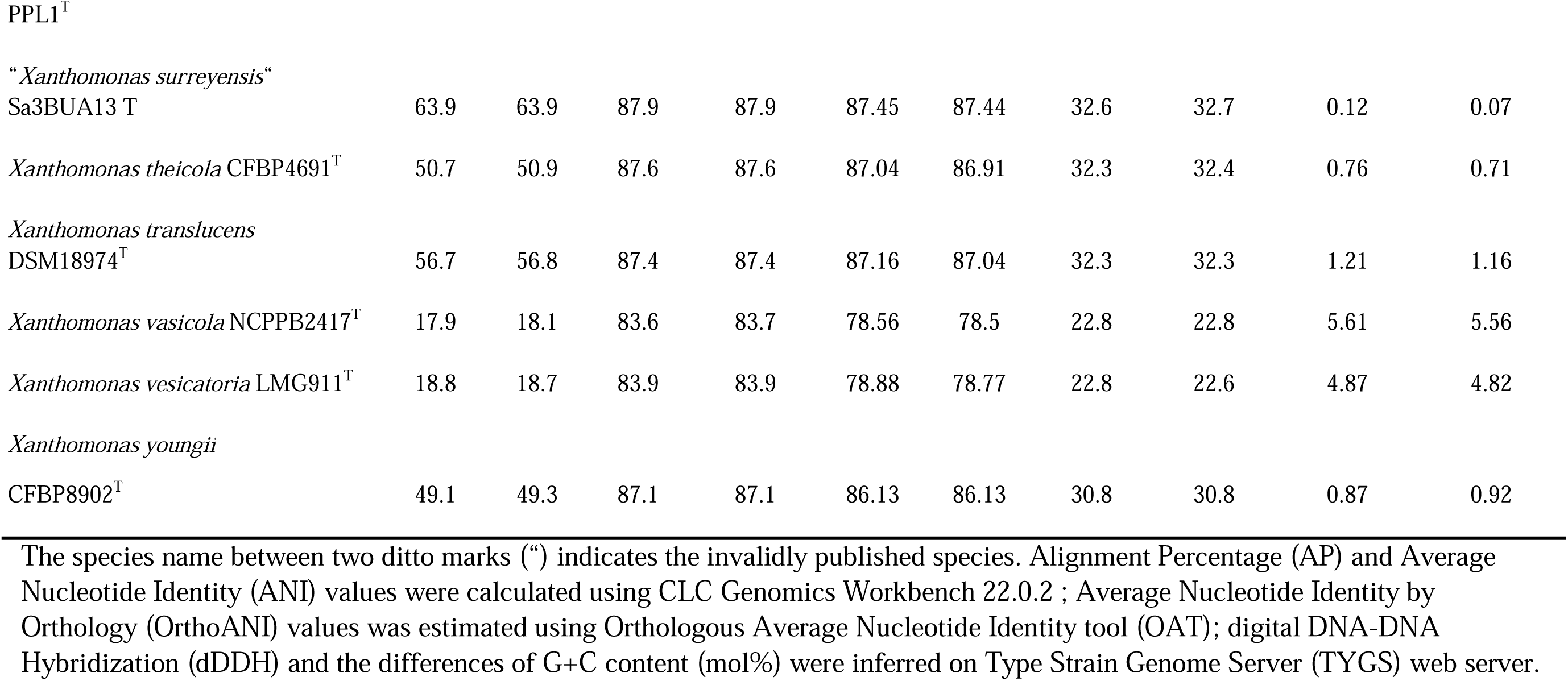
Overall genomic relatedness indices (OGRIs) comparison of new species, *Xanthomonas hawaiienesis* sp. nov., strains with other type strains of *Xanthomonas* species.

The estimations of ANI and dDDH of anthurium strains, A5588^T^ and A5586^T^, were 86.4% and 28.2%, respectively. Both strains shared ANI and dDDH values lower than 90% (83.4-86.8%) and 30% (20.7%-29.9%) with other type strains of *Stenotrophomonas* spp., respectively, except A5588^T^ and *S. bentonitica* LMG 29893^T^ shared 94.7% of ANI and 56.4% of dDDH sequence identities (Table 2). Besides ANI and dDDH values, other OGRIs including AP, OrthoANI, and G+C differences supported A5588^T^ and A5586^T^ are two novel species (Table 3). To combine the phylogenetic analyses and OGRIs evidence, three novel species were proposed, i.e. *X. hawaiiensis* sp. nov. strains A6251^T^ and A2111; *S. aracearum* sp. nov. strain A5588^T^ and *S. oahuensis* sp. nov. strain A5586^T^.

**TABLE 3.**
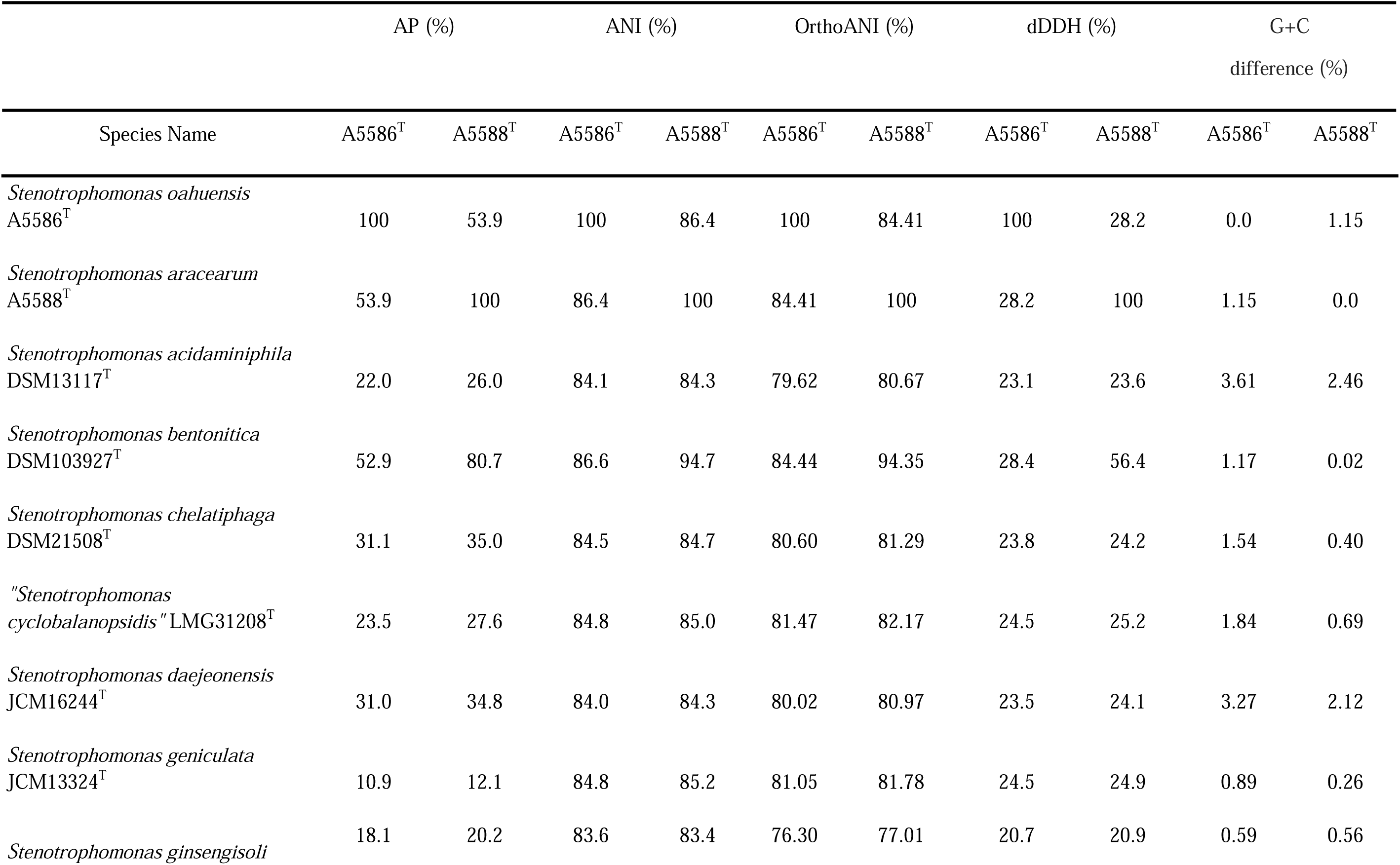
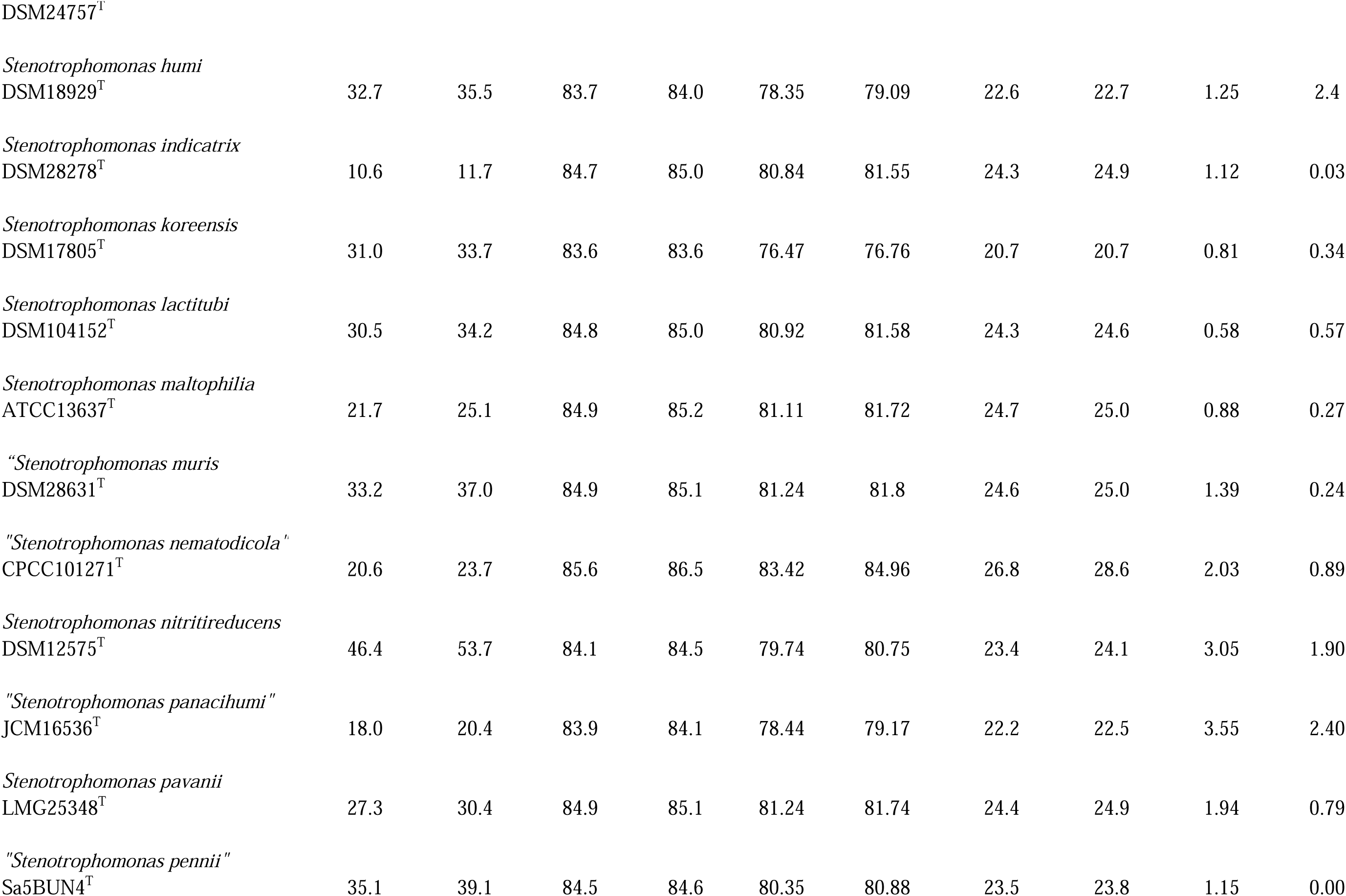
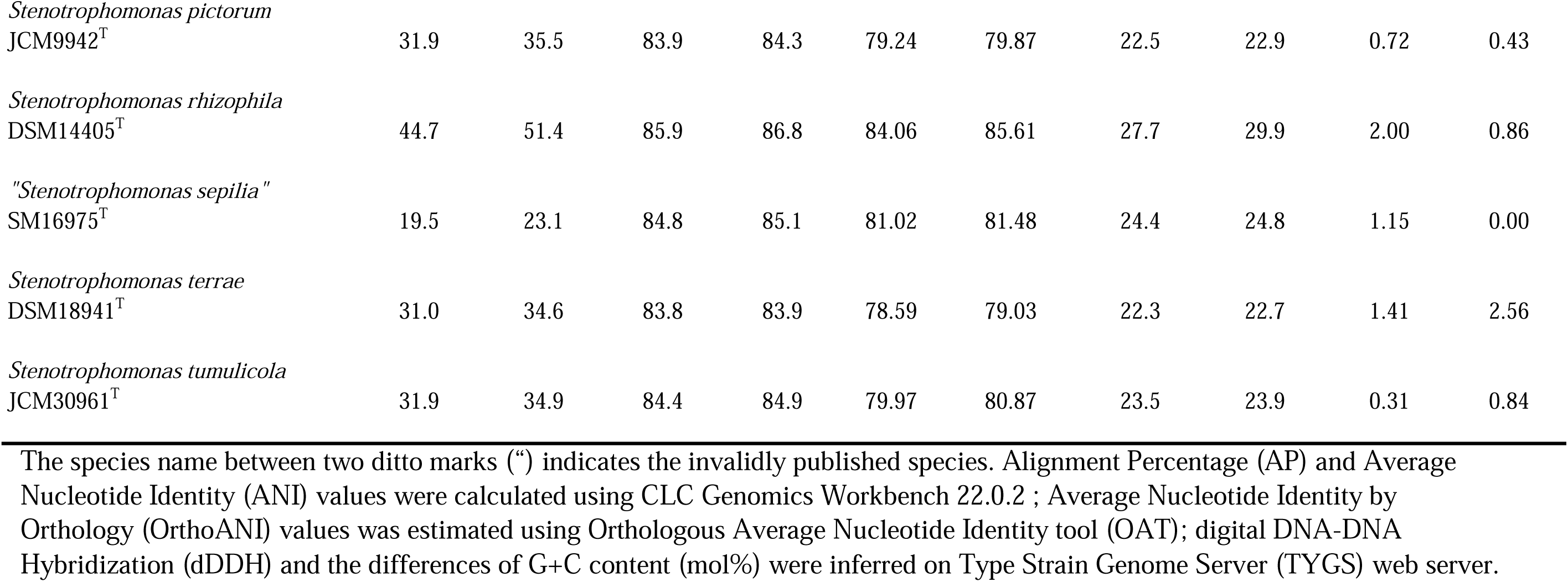
Overall genomic relatedness indices (OGRIs) comparison of new species, *Stenotrophomonas oahuensis* sp. nov., and *S. aracearum* sp. nov. with other species in the genus.

| Species Name | AP (%) |  | ANI (%) |  | OrthoANI (%) |  | dDDH (%) |  | G+C difference (%) |  |
| --- | --- | --- | --- | --- | --- | --- | --- | --- | --- | --- |
|  | A5586<sup>T</sup> | A5588<sup>T</sup> | A5586<sup>T</sup> | A5588<sup>T</sup> | A5586<sup>T</sup> | A5588<sup>T</sup> | A5586<sup>T</sup> | A5588<sup>T</sup> | A5586<sup>T</sup> | A5588<sup>T</sup> |
| *Stenotrophomonas oahuensis*<br>A5586<sup>T</sup> | 100 | 53.9 | 100 | 86.4 | 100 | 84.41 | 100 | 28.2 | 0.0 | 1.15 |
| *Stenotrophomonas aracearum*<br>A5588<sup>T</sup> | 53.9 | 100 | 86.4 | 100 | 84.41 | 100 | 28.2 | 100 | 1.15 | 0.0 |
| *Stenotrophomonas acidaminiphila*<br>DSM13117<sup>T</sup> | 22.0 | 26.0 | 84.1 | 84.3 | 79.62 | 80.67 | 23.1 | 23.6 | 3.61 | 2.46 |
| *Stenotrophomonas bentonitica*<br>DSM103927<sup>T</sup> | 52.9 | 80.7 | 86.6 | 94.7 | 84.44 | 94.35 | 28.4 | 56.4 | 1.17 | 0.02 |
| *Stenotrophomonas chelatiphaga*<br>DSM21508<sup>T</sup> | 31.1 | 35.0 | 84.5 | 84.7 | 80.60 | 81.29 | 23.8 | 24.2 | 1.54 | 0.40 |
| "*Stenotrophomonas*<br>*cyclobalanopsidis*" LMG31208<sup>T</sup> | 23.5 | 27.6 | 84.8 | 85.0 | 81.47 | 82.17 | 24.5 | 25.2 | 1.84 | 0.69 |
| *Stenotrophomonas daejeonensis*<br>JCM16244<sup>T</sup> | 31.0 | 34.8 | 84.0 | 84.3 | 80.02 | 80.97 | 23.5 | 24.1 | 3.27 | 2.12 |
| *Stenotrophomonas geniculata*<br>JCM13324<sup>T</sup> | 10.9 | 12.1 | 84.8 | 85.2 | 81.05 | 81.78 | 24.5 | 24.9 | 0.89 | 0.26 |
| *Stenotrophomonas ginsengisoli* | 18.1 | 20.2 | 83.6 | 83.4 | 76.30 | 77.01 | 20.7 | 20.9 | 0.59 | 0.56 |
| DSM24757<sup>T</sup> |  |  |  |  |  |  |  |  |  |  |
| *Stenotrophomonas humi*<br>DSM18929<sup>T</sup> | 32.7 | 35.5 | 83.7 | 84.0 | 78.35 | 79.09 | 22.6 | 22.7 | 1.25 | 2.4 |
| *Stenotrophomonas indicatrix*<br>DSM28278<sup>T</sup> | 10.6 | 11.7 | 84.7 | 85.0 | 80.84 | 81.55 | 24.3 | 24.9 | 1.12 | 0.03 |
| *Stenotrophomonas koreensis*<br>DSM17805<sup>T</sup> | 31.0 | 33.7 | 83.6 | 83.6 | 76.47 | 76.76 | 20.7 | 20.7 | 0.81 | 0.34 |
| *Stenotrophomonas lactitubi*<br>DSM104152<sup>T</sup> | 30.5 | 34.2 | 84.8 | 85.0 | 80.92 | 81.58 | 24.3 | 24.6 | 0.58 | 0.57 |
| *Stenotrophomonas maltophilia*<br>ATCC13637<sup>T</sup> | 21.7 | 25.1 | 84.9 | 85.2 | 81.11 | 81.72 | 24.7 | 25.0 | 0.88 | 0.27 |
| "*Stenotrophomonas muris*<br>DSM28631<sup>T</sup> | 33.2 | 37.0 | 84.9 | 85.1 | 81.24 | 81.8 | 24.6 | 25.0 | 1.39 | 0.24 |
| "*Stenotrophomonas nematodicola*"<br>CPCC101271<sup>T</sup> | 20.6 | 23.7 | 85.6 | 86.5 | 83.42 | 84.96 | 26.8 | 28.6 | 2.03 | 0.89 |
| *Stenotrophomonas nitritireducens*<br>DSM12575<sup>T</sup> | 46.4 | 53.7 | 84.1 | 84.5 | 79.74 | 80.75 | 23.4 | 24.1 | 3.05 | 1.90 |
| "*Stenotrophomonas panacihumi*"<br>JCM16536<sup>T</sup> | 18.0 | 20.4 | 83.9 | 84.1 | 78.44 | 79.17 | 22.2 | 22.5 | 3.55 | 2.40 |
| *Stenotrophomonas pavanii*<br>LMG25348<sup>T</sup> | 27.3 | 30.4 | 84.9 | 85.1 | 81.24 | 81.74 | 24.4 | 24.9 | 1.94 | 0.79 |
| "*Stenotrophomonas pennii*"<br>Sa5BUN4<sup>T</sup> | 35.1 | 39.1 | 84.5 | 84.6 | 80.35 | 80.88 | 23.5 | 23.8 | 1.15 | 0.00 |
| *Stenotrophomonas pictorum*<br>JCM9942<sup>T</sup> | 31.9 | 35.5 | 83.9 | 84.3 | 79.24 | 79.87 | 22.5 | 22.9 | 0.72 | 0.43 |
| *Stenotrophomonas rhizophila*<br>DSM14405<sup>T</sup> | 44.7 | 51.4 | 85.9 | 86.8 | 84.06 | 85.61 | 27.7 | 29.9 | 2.00 | 0.86 |
| "*Stenotrophomonas seipilia*"<br>SM16975<sup>T</sup> | 19.5 | 23.1 | 84.8 | 85.1 | 81.02 | 81.48 | 24.4 | 24.8 | 1.15 | 0.00 |
| *Stenotrophomonas terrae*<br>DSM18941<sup>T</sup> | 31.0 | 34.6 | 83.8 | 83.9 | 78.59 | 79.03 | 22.3 | 22.7 | 1.41 | 2.56 |
| *Stenotrophomonas tumulicola*<br>JCM30961<sup>T</sup> | 31.9 | 34.9 | 84.4 | 84.9 | 79.97 | 80.87 | 23.5 | 23.9 | 0.31 | 0.84 |

The species name between two ditto marks ("") indicates the invalidly published species. Alignment Percentage (AP) and Average Nucleotide Identity (ANI) values were calculated using CLC Genomics Workbench 22.0.2 ; Average Nucleotide Identity by Orthology (OrthoANI) values was estimated using Orthologous Average Nucleotide Identity tool (OAT); digital DNA-DNA Hybridization (dDDH) and the differences of G+C content (mol%) were inferred on Type Strain Genome Server (TYGS) web server.

### Pan– and Core-Genomic Analyses

Among 40 reference genomes of *Xanthomonas* spp. including *X. hawaiiensis* sp. nov. strains A6251^T^ and A2111, total 425 core orthologous genes (99%≤ strains ≤ 100%) and 28,285 cloud genes (0%≤ strains < 15%) were found (Figure 4a). The lowest two numbers of unique genes present in A6251^T^ and A2111 were 87 and 103, respectively, and following *X. sacchari* CFBP4641^T^ had 171 unique genes in the Figure 4b. The number of exclusive hypothetical protein encoded genes were comparatively lower in strains A6251^T^ and A2111, whereas 50 common hypothetical proteins existed in all type strains of *Xanthomonas* spp. (Figure 4c). Based on the phylogenetic tree constituted with 425 core genes, the closest relative of *X. hawaiiensis* sp. nov. was *X. sacchari* CFBP4641^T^ and successively clustered with *“X. sontii”* PPL1^T^ and “*X. indica*” CFBP9039^T^ in *Xanthomonas* clade I species (Figure 5). The groupings were concordant with previously described MLSA tree (Figure 3). The 34,713 gene clusters estimated in the Roary matrix revealed that xanthomonads’ genomes were highly diversified (Figure 5).

**Figure 4.**
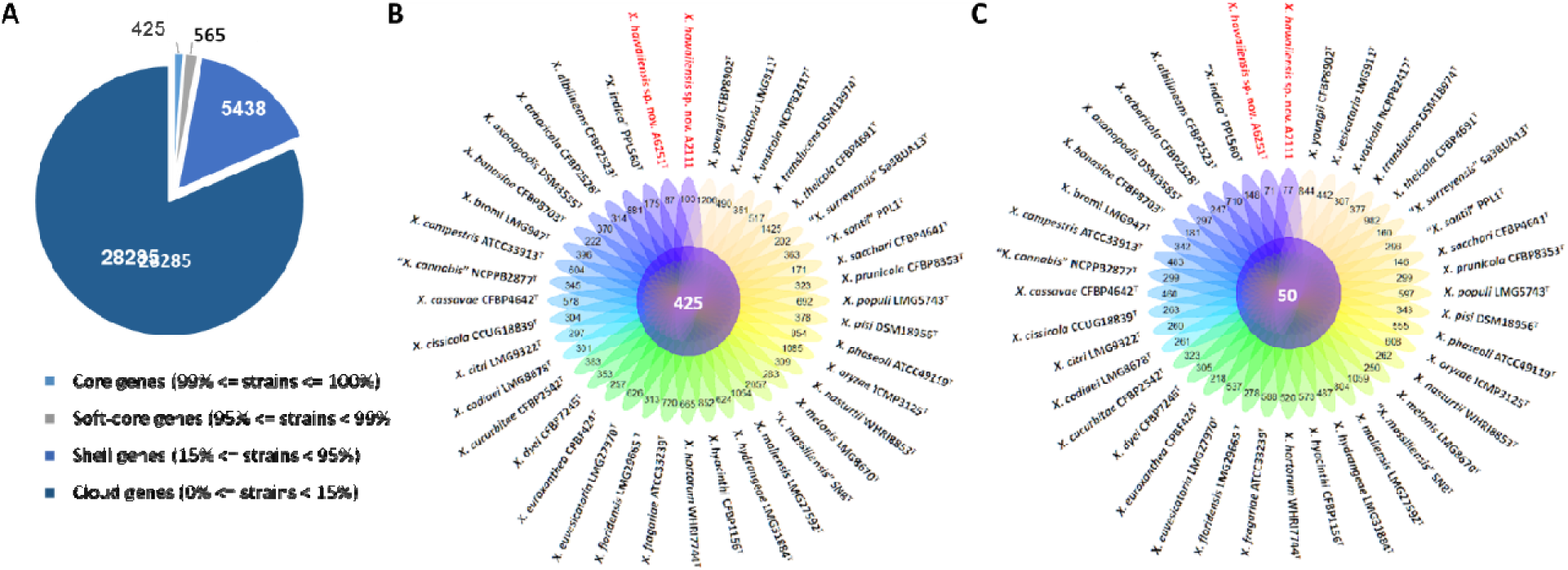
Pan-genome analyses of *Xanthomonas hawaiienses* sp. nov. (A6251^T^ and A2111) with other type strains of *Xanthomonas* species. (A) Numbers of core, soft-core, shell, and cloud genes within 40 genomes of type strains of *Xanthomonas* species. (B) Floral plot showing the number of core orthologous genes in the center and the number of unique genes on each petal. (C) The number of common hypothetical protein encoding genes in the center of floral plot and the number of unique hypothetical protein encoding genes of each *Xanthomonas* strain on each petal.

**Figure 5.**
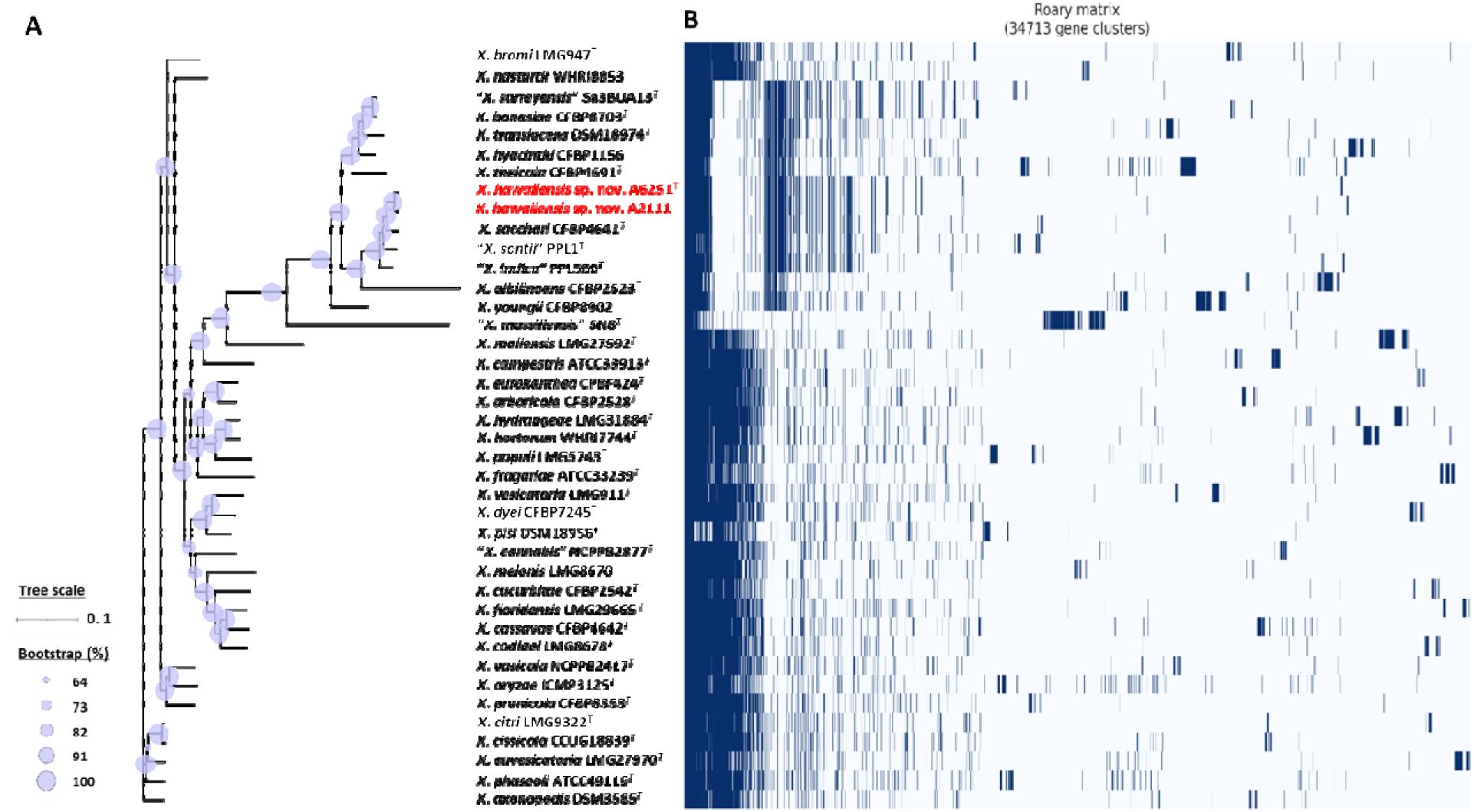
Core and pan genome analyses of 40 *Xanthomonas* species including new species strains. (A) Core genome-based ML phylogenetic tree of *Xanthomonas hawaiienses* sp. nov. (A6251^T^ and A2111) with other type strains of *Xanthomonas* species. Tree scale bar represents the nucleotide substitutions per site. The range of purple circles indicate the percentage of bootstrapping confidence. (B) Pan genome-based Roary matrix of present and absent genes among 40 coordinated *Xanthomonas* species. Dark blue blocks present genes and pale blue blocks are missing genes in the genomes.

On the other hand, the pan genome size of 20 *Stenotrophomonas* spp. type strains, including *S. aracearum* sp. nov. (A5588^T^) and *S. oahuensis* sp. nov. (A5586^T^), was 31,069 with 576 core genes (Figure 6a, 7). The genome of strain A5588^T^ contained 396 unique genes and 317 of which were hypothetical protein encoding genes; whereas, high number of hypothetical protein encoding genes (1,242 genes) were harbored in the genome of strain A5586^T^ possessing total 1,526 unique genes (Figure 6b-c). As presented in 9-gene ML tree (Figure 3), A5588^T^ and *S. bentonitica* DSM103927^T^ were closely clustered together and grouped with A5586^T^, which was a sister group of the clade formed with *S. rhizophila* DSM14405^T^ and *“S. nematodicola”* CPCC101271 (Figure 7). The average number of unique genes with unknown functions was higher in 25 *Stenotrophomonas* spp. than 40 *Xanthomonas* spp. (806 > 538), implying higher genetic diversity within stenotrophomonads, warrant further investigations on *Stenotrophomonas* species.

**Figure 6.**
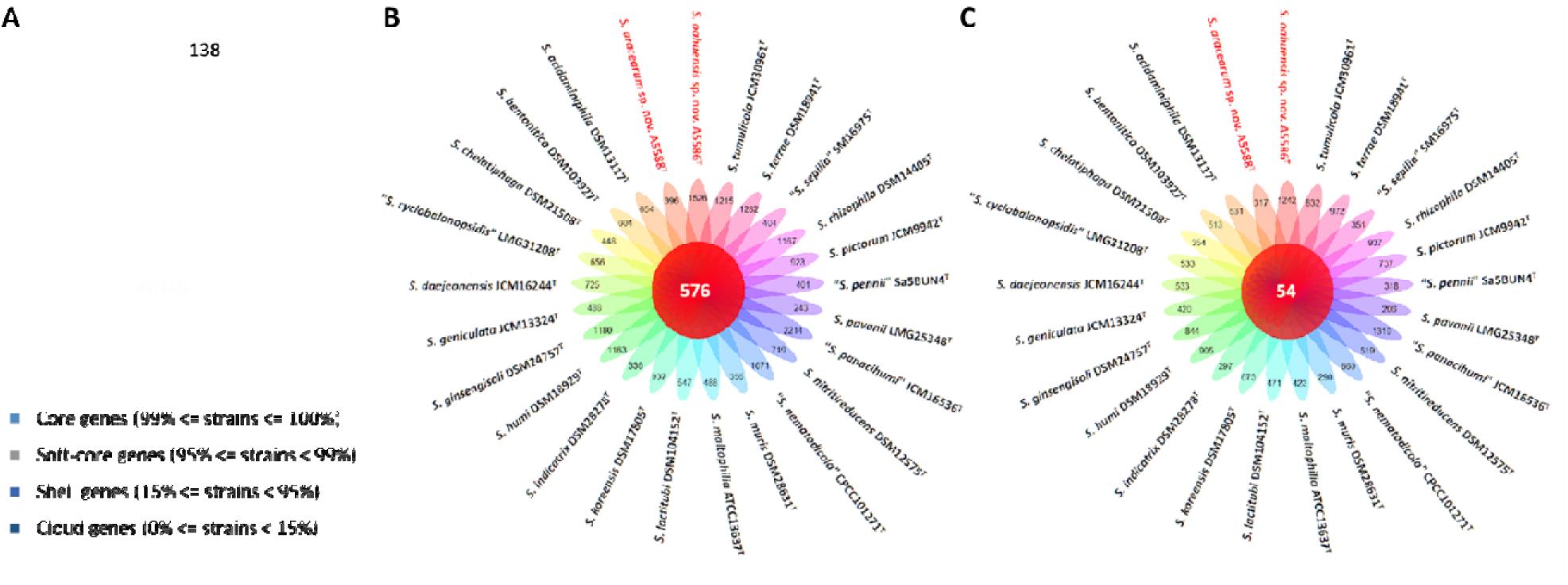
Pan-genome analyses of *Stenotrophomonas aracearum* sp. nov. (A5588^T^) and *S. oahuensis* sp. nov. (A5586^T^) with other type strains of *Stenotrophomonas* species. (A) Numbers of core, soft-core, shell, and cloud genes within 25 genomes of type strains of *Stenotrophomonas* species. (B) Floral plot showing the number of core orthologous genes in the center and the number of unique genes on each petal. (C) The number of common hypothetical protein encoding genes in the center of floral plot and the number of unique hypothetical protein encoding genes of each *Stenotrophomonas* species on each petal.

**Figure 7.**
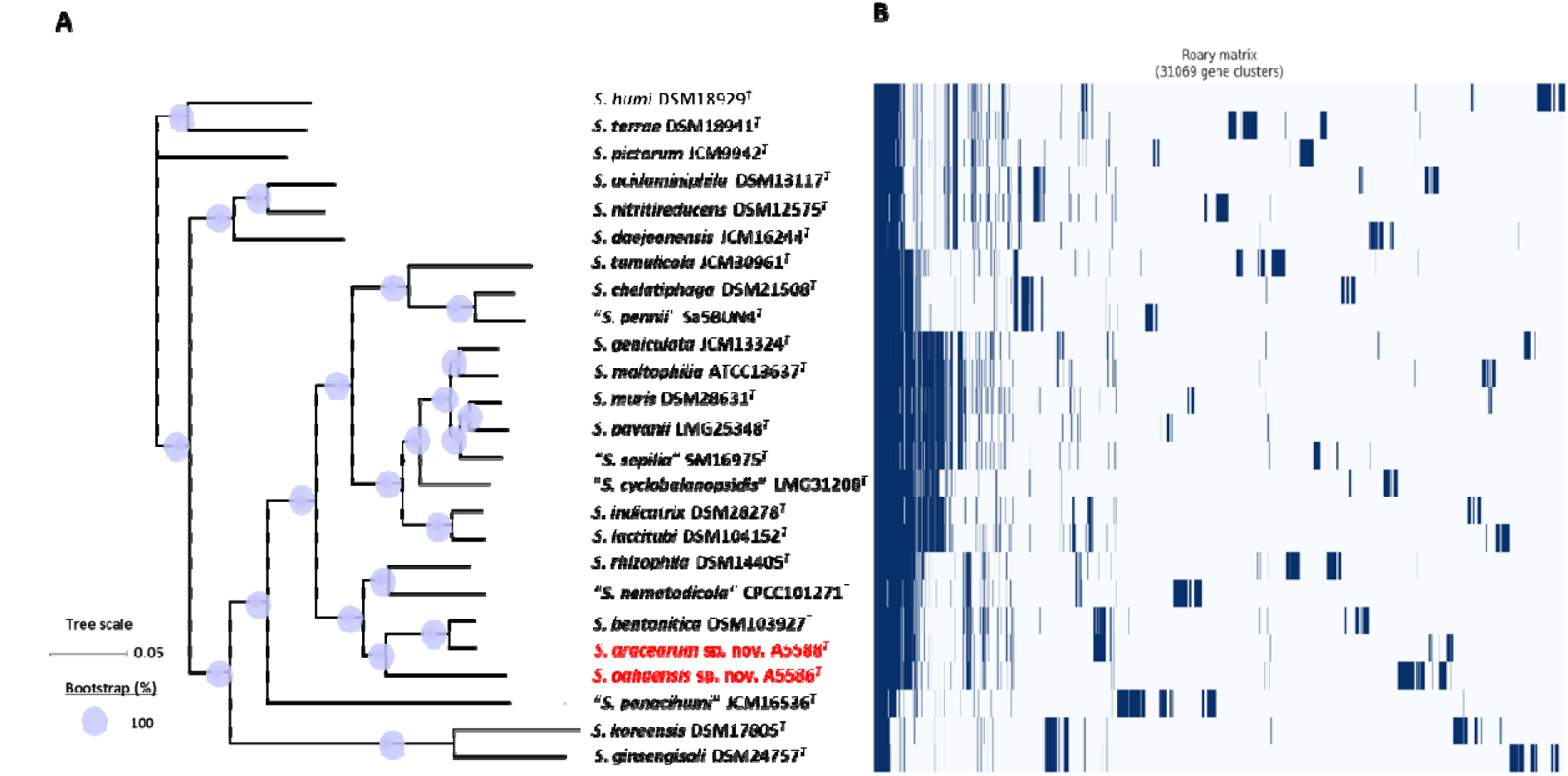
Core and pan genome analyses of 25 *Stenotrophomonas* species including two new species type strains. (A) Core genome-based ML phylogenetic tree of *S. aracearum* sp. nov. (A5588^T^) and *S. oahuensis* sp. nov. (A5586^T^) along with other type strains of *Stenotrophomonas* species. Tree scale bar represents the substitutions per nucleotide position. The purple circles represent 100% bootstrapping support. (B) Pan genome based Roary matrix of present and absent genes among all coordinated *Stenotrophomonas* species. Dark blue blocks indicate genes present and pale blue blocks indicate genes absent in the genomes.

### Antibiotic Sensitivity Assays

The inhibition zones with seven tested antibiotics, bacitracin (50 mg/ml), chloramphenicol (50 mg/ml), gentamicin (50 mg/ml), kanamycin (50 mg/ml), penicillin (50 mg/ml), tetracycline (40 mg/ml), and polymyxin B sulfate (50 mg/ml), indicated various degrees of sensitivity of four new species strains. Strains A6251^T^, A2111, and A5586^T^ were sensitive to all tested antibiotics, whereas strain A5588^T^ was sensitive to all tested antibiotics except penicillin (Table S2). Strains A6251^T^ and A2111, belonging to the same new species, displayed similar results except the tolerance to polymyxin B sulfate. Notably, A5586^T^ displayed a very small inhibition zone (0.1 cm in radius) surrounding the discs of bacitracin on the NA plate after incubating at 28 °C for 24 hours (Table S2).

### Descriptions of New Species

***Xanthomonas hawaiiensis* sp. nov.** (ha.wai.i.en′sis N.L. masc./fem. adj. *hawaiiensis* of or belonging to Hawaii, a state of the United State, referring to the geographical origin of the new species).

Colonies of the type strain A6251^T^ are yellow (Honey, Hex code #FFC30B), circular shape, mucoid consistency, smooth surface, convex relief with entire margins, and 0.3-0.6 (avg. 0.45) mm in diameter on yeast dextrose calcium carbonate (YDC) medium plate after incubating at 28 °C for 2 days. Cells are Gram-negative and able to utilize Dextrin, D-Maltose, D-Trehalose, D-Cellobiose, Gentiobiose, Sucrose, D-Turanose, α– D-Lactose, D-Melibiose, Β-Methyl-D-Glucoside, D-Salicin, N-Acetyl-D-Glucosamine, α– D-Glucose, D-Mannose, D-Fructose, D-Galactose, L-Fucose, 1% NaCl, 1% Sodium Lactate, Glycerol, Gelatin, L-Glutamic Acid, Lincomycin, Pectin, Quinic Acid, Vancomycin, Tetrazolium Violet, Tetrazolium Blue, Citric Acid, Bromo-Succinic Acid, Lithium Chloride, Tween 40, and Acetic Acid. In contrast, cells are unable to oxidate Stachyose, D-Raffinose, N-Acetyl-β-D-Mannosamine, N-Acetyl-D-Galactosamine, N-Acetyl-Neuraminic Acid, 8% NaCl, Inosine, Fusidic Acid, D-Sorbitol, D-Mannitol, D-Arabitol, myo-Inositol, D-Aspartic Acid, Minocycline, L-Arginine, L-Histidine, L-Pyroglutamic Acid, Guanidine HCl, D-Gluconic Acid, Mucic Acid, D-Saccharic Acid, p-Hydroxy-Phenylacetic Acid, D-Lactic Acid Methyl Ester, α-Keto-Glutaric Acid, D-Malic Acid, γ-Amino-Butryric Acid, α-Hydroxy-Butyric Acid, α-Keto-Butyric Acid, Formic Acid, Sodium Butyrate, and Sodium Bromate. Some utilizations of carbon resources and chemical components showed borderline results or inconsistency between two strains after growing cell suspension in GEN III Microplate (Biolog Inc., Hayward, CA) at 28 °C for 24 hours.

*X. hawaiiensis* sp. nov. was sensitive to seven tested antibiotics, including Bacitracin (50 mg/ml), Chloramphenicol (50 mg/ml), Gentamicin (50 mg/ml), Kanamycin (50 mg/ml), Penicillin (50 mg/ml), Tetracycline (40 mg/ml), and Polymyxin B Sulfate (50 mg/ml). The genome size of type strain A6251^T^ is 4.88 Mbp with 68.93 mol% of DNA G+C content.

The type strain A6251^T^ =D-93^T^ =ICMP25022^T^ =LMG33200^T^ was isolated from *Spathiphyllum* (Araceae family) in 1985 in Hawaii, USA. Another strain A2111 =D-194 =ICMP25023 =LMG33199 was isolated from *Colocasia* (Araceae family) in 1986 in Hawaii, USA.

***Stenotrophomonas aracearum* sp. nov.** (a.ra.ce.a’rum. N.L. gen. fem. pl. n. aracearum, representatives of plants belonging to the Araceae family).

Colonies of *S. aracearum* strain A5588^T^ are dark yellow (Mustard, Hex code #E8B828), irregular shape, butyrous consistency, smooth surface, raised relief with entire margins, and 0.4-0.5 (ave. 0.45) mm in diameter on YDC medium plate after incubation at 28 °C for 2 days. Cells are Gram-negative and able to utilize D-maltose, D-cellobiose, gentiobiose, N-acetyl-D-glucosamine, N-acetyl-D-galactosamine, α-D-glucose, D-mannose, 1% sodium lactate, D-serine, troleandomycin, rifamycin SV, gelatin, lincomycin, guanidine HCl, vancomycin, tetrazolium violet, tetrazolium Blue, α-keto-glutaric acid, L-malic acid, bromo-succinic acid, acetic acid, and aztreonam. Cells grow under pH6 and 1% NaCl but not at pH5 nor 8% NaCl. In the contrary, cells are unable to oxidize sucrose, D-turanose, stachyose, D-raffinose, α-D-lactose, D-melibiose, Β-methyl-D-glucoside, N-acetyl-β-D-mannosamine, N-acetyl-neuraminic acid, D-galactose, 3-methyl-glucose, inosine, fusidic acid, D-sorbitol, D-mannitol, D-arabitol, myo-inositol, glycerol, D-glucose-6-PO4, D-aspartic acid, D-serine, minocycline, L-arginine, L-aspartic acid, L-glutamic acid, L-histidine, L-pyroglutamic acid, L-serine, pectin, D-galacturonic acid, D-gluconic acid, mucic acid, quinic acid, D-saccharic acid, p-hydroxy-phenylacetic acid, D-lactic acid methyl ester, L-lactic acid, citric acid, D-malic acid, nalidixic acid, potassium tellurite, γ-amino-butryric acid, α-hydroxy-butyric acid, α-keto-butyric acid, β-hydroxy-D, L-butyric acid, acetoacetic acid, formic acid, sodium butyrate, and sodium bromate. Some utilizations of carbon sources and chemical components, such as dextrin and glucuronamide showed faded positive results after growing A5588^T^ cell suspension in GEN III Microplate (Biolog Inc., Hayward, CA) at 28 °C for 24 hours.

*S. aracearum* sp. nov. was sensitive to six tested antibiotics including Bacitracin (50 mg/ml), Chloramphenicol (50 mg/ml), Gentamicin (50 mg/ml), Kanamycin (50 mg/ml), Tetracycline (40 mg/ml), and Polymyxin B Sulfate (50 mg/ml) but resistant to Penicillin (50 mg/ml) on NA plates. The genome size of type strain A5588^T^ is of 4.33 Mbp with 66.44 mol% of DNA G+C content.

The type strain A5588^T^ =D-61-1L^T^ =ICMP25025^T^ =LMG33202^T^ was isolated from *Anthurium* (Araceae family) in 1985 in Hawaii, USA.

***Stenotrophomonas oahuensis* sp. nov.** (o.a.hu.en’sis N.L. masc./fem. adj. *oahuensis* of or belonging to the island of Oahu in Hawaii, referring to the geographical origin of the new species).

Colonies of the *S. oahuensis* strain A5586^T^ are dark yellow (Butterscotch, Hex code #FABD02), circular shape, butyrous consistency, smooth surface, flat relief with undulate margins, and 0.4-0.7 (ave. 0.55) mm in diameter on YDC medium plate after incubation at 28 °C for 2 days. Cells are Gram-negative and able to utilize dextrin, D-maltose, D-trehalose, D-cellobiose, gentiobiose, Β-methyl-D-glucoside, D-salicin, N-acetyl-D-glucosamine, α-D-glucose, D-mannose, 1% sodium lactate, gelatin, glycyl-L-proline, lincomycin, guanidine HCl, vancomycin, tetrazolium violet, tetrazolium Blue, citric acid, α-keto-glutaric acid, L-malic acid, bromo-succinic acid, lithium chloride, propionic acid, acetic acid, and aztreonam. Cells grow under the conditions of pH 6, 1% NaCl or 4% NaCl but cells survive neither pH 5 nor 8% NaCl solution. In contrast, cells are unable to oxidize sucrose, stachyose, D-raffinose, N-acetyl-β-D-mannosamine, N-acetyl-neuraminic acid, D-galactose, 3-methyl-glucose, inosine, D-fucose, L-fucose, L-rhamnose, inosine, fusidic acid, D-sorbitol, D-mannitol, D-arabitol, myo-inositol, glycerol, D-glucose-6-PO4, D-aspartic acid, D-serine, rifamycin SV, minocycline, L-arginine, L-aspartic acid, L-glutamic acid, L-histidine, L-pyroglutamic acid, L-serine, pectin, D-galacturonic acid, D-gluconic acid, D-glucuronic acid, mucic acid, quinic acid, D-saccharic acid, p-hydroxy-phenylacetic acid, D-lactic acid methyl ester, L-lactic acid, D-malic acid, nalidixic acid, potassium tellurite, γ-amino-butryric acid, α-hydroxy-butyric acid, β-hydroxy-D, L-butyric acid, α-keto-butyric acid, acetoacetic acid, formic acid, and sodium bromate. Some utilizations of carbon resources and chemical components, such as D-turanose and sodium butyrate showed faded positive results after growing A5586^T^ cell suspension in GEN III Microplate (Biolog Inc., Hayward, CA) at 28 °C for 24 hours.

*S. oahuensis* sp. nov. was sensitive to seven tested antibiotics including Bacitracin (50 mg/ml), Chloramphenicol (50 mg/ml), Gentamicin (50 mg/ml), Kanamycin (50 mg/ml), Penicillin (50 mg/ml), Tetracycline (40 mg/ml), and Polymyxin B Sulfate (50 mg/ml). The genome size of type strain A5586^T^ is 4.68 Mbp, which includes a chromosome (4.62 Mbp) and a plasmid (60.78 Kbp). The DNA G+C content of the type strain is 65.3 mol%. The type strain A5586^T^ =D-31^T^ =ICMP25024^T^ =LMG33201^T^ was isolated from *Anthurium* (Araceae family) in 1981 in Hawaii, USA.

## Discussion

The genera *Xanthomonas* and *Stenotrophomonas* are phylogenetically and evolutionary linked and also found frequently together in several niches, including environmental reservoirs (plants and soil) and biofilters used for waste gas treatment of animal-rendering plant (Lipski and Altendorf, 1997; Finkmann et al. 2000; Ryan et al. 2009).

Although more studies are focused on phyto– and human-pathogenic species, the versatility of *Xanthomonas* and *Stenotrophomonas* spp. have potential to apply to many different fields and needed to be explored further.

The well-known industrial biopolymer and food additive is xanthan gum produced from *X. campestris* and other *Xanthomonas* species (Margaritis and Zajic, 1978; Kennedy and Bradshaw, 1984; Gumus et al. 2010). Other productions of bioactive secondary metabolites from xanthomonads are pigment xanthomonadin, analogues of which have antioxidant potential, and xanthoferrin, which acts as a bioproduction agent under low iron conditions (Madden et al. 2019; Pandey et al. 2017). The increasing number of studies on non-pathogenic xanthomonads isolated from rice, banana, citrus, walnut and so on suggest they have the potential of biocontrol and bioprotection against the causal agent of their host plants (Bansal et al. 2021; Rana et al. 2022; Fernandes et al. 2021). For example, *X. sontii* strain R1 (formerly misclassified as *X. sacchari*) isolated from rice seed was reported to have an antagonistic ability against *Burkholderia glumae*, which causes rice panicle blight disease (Xie et al. 2003; Ham et al. 2011; Fang et al. 2015). In addition, *Xanthomonas* sp. from ryegrass, which was phylogenetically closely related to *X. translucens*, showed bioprotection activities against broad tested fungal pathogens (Li et al. 2020).

Recently, more research on the potentials of biotechnological applications and biological controls of stenotrophomonads were unraveled. In agriculture, for example, *Stenotrophomonas* strains were known for promoting plant growth, protecting plant against biotic and abiotic stresses, and treating as biocontrol for plant diseases (Wolf et al. 2002; Alavi et al. 2013; Berg et al. 2015; Zhang and Yuen, 1999; Messiha et al. 2007). As bioremediators and phytoremediators, *Stenotrophomonas* strains are capable of metabolizing and degrading a broad range of organic compounds, such as benzene and toluene, and tolerating antibiotics and heavy metals, such as mercury and silver (Binks et al. 1995; Lee et al. 2002; Alonso et al. 2000; Pages et al. 2008).

In this study, we proposed three new species, *X. hawaiiensis* sp. nov., *S. aracearum* sp. nov. and *S. oahuensis* sp. nov. isolated from Araceae. Interestingly, the high numbers of unique genes and hypothetical protein encoding genes unraveled from detailed genomic contents of the novel species, especially in *S. oahuensis* sp. nov., imply that more novel or useful enzymatic properties and metabolic capabilities of *Xanthomonas* and *Stenotrophomonas* spp. from different environmental sources are worth exploring for biocontrol and bioprotection purposes. Preliminary data from pathogenicity tests on anthurium indicated that strains A5588^T^ and A5586^T^ from anthurium are non-pathogenic stenotrophomonads due to no symptom development on their original host. Hence, further studies of their pathogenesis on Araceae and other possible bioactivities should be performed.

## Acknowledgements

This research was funded by the USDA National Institute of Food and Agriculture, Hatch project 9038H, managed by the College of Tropical Agriculture and Human Resources. This work was also supported by the USDA-ARS Agreement No. 58-2040-9-011, Systems Approaches to Improve Production and Quality of Specialty Crops Grown in the U.S. Pacific Basin; sub-project: Genome Informed Next Generation Detection Protocols for Pests and Pathogens of Specialty Crops in Hawaii. The strains were stored and maintained by the National Science Foundation funded project (NSF-CSBR Grant No. DBI-1561663). The bioinformatics analyses were supported by NIGMS of the National Institutes of Health under award number P20GM125508.

