## Supplement Tables for "Three new species, *Xanthomonas hawaiiensis* sp. nov., *Stenotrophomonas aracearum* sp. nov., and *Stenotrophomonas oahuensis* sp. nov., isolated from Araceae family": Supplement Tables.pdf

**TABLE S1**. The strain information and genome accession numbers of four new species strains and the type strains of *Xanthomonas* and *Stenotrophomonas* spp. analyzed in this study.

| Species Name | Type Strain Number | Isolate Source | Geographic Origin | Year | Assembly Accession Number | Reference |
|---|---|---|---|---|---|---|
| ***Stenotrophomonas* genus** | | | | | | |
| *Stenotrophomonas aracearum* sp. nov. | A5588; D-61-1L | *Anthurium* | USA: Hawaii | 1985 | CP115541-CP115542 | This study |
| *Stenotrophomonas oahuensis* sp. nov. | A5586; D-31 | *Anthurium* | USA: Hawaii | 1981 | CP115543 | This study |
| *Stenotrophomonas acidaminiphila* | AMX 19; DSM 13117 | Anaerobic sludge | Mexico | 2002 | GCF_024221815.1 | Assih et al. 2002 |
| *Stenotrophomonas bentonitica* | BII-R7; CECT 9180; DSM 103927; LMG 29893 | Bentonite formations | China | 2019 | GCF_013185915.1 | Sánchez-Castro et al. 2017 |
| *Stenotrophomonas chelatiphaga* | CCUG 57178; DSM 21508; LPM-5; VKM B-2486 | Municipal sewage sludge | Russia | 2009 | GCF_001431535.1 | Kaparullina et al. 2010 |
| *Stenotrophomonas daejeonensis* | DSM 26149; JCM 16244; KCTC 22451; MJ03 | Sewage water | South Korea | 2010 | GCF_001431505.1 | Lee et al. 2011 |
| *Stenotrophomonas geniculata* | ATCC 19374; JCM 13324; LMG 2195; NCIB 9428; NCIMB 9428 | Tap water | USA | - | GCF_001431625.1 | (Wright 1895) Rudra and Gupta 2021 |
| *Stenotrophomonas ginsengisoli* | DCY 1; DSM 24757; KCTC 12539; NBRC 101154 | Soil from ginseng field | South Korea | 2010 | GCF_001431485.1 | Kim et al. 2010 |
| *Stenotrophomonas humi* | DSM 18929; LMG 23959; R-32729 | Soil | Belgium | 2007 | GCF_001431415.1 | Heylen et al. 2007 |
| *Stenotrophomonas indicatrix* | DSM 28278; LMG 29942; WS40 | Dirty dishes | Germany | 2013 | GCF_002750975.1 | Weber et al. 2018 |
| *Stenotrophomonas koreensis* | DSM 17805; JCM 13256; KCTC 12211; TR6-01 | Compost | South Korea | 2003 | GCF_001431525.1 | Yang et al. 2006 |
| *Stenotrophomonas lactitubi* | DSM 104152; LMG 29943; | Milking machine | Germany | 2014 | GCF_002803515.1 | Weber et al. |

| | | | | | | |
|---|---|---|---|---|---|---|
| | M15 | biofilm | | | | 2018 |
| *Stenotrophomonas maltophilia* | ATCC 13637 ;DSM 50170 ;IFO 14161 ;ICPB 2648-67 ;NCIB 9203 ;NCPPB 1974 ;ICMP 17033 | Oropharyngeal region of patient with mouth cancer | USA | 1959 | GCF_001997185.1 | (Hugh 1981) Palleroni and Bradbury 1993 |
| *Stenotrophomonas nitritireducens* | ATCC BAA-12; CCUG 46888; CIP 107228; DSM 12575; JCM 13311; L2 | Laboratory scale biofilters supplied with ammonia or dimethyl disulfide and ammonia | Germany | 1997 | GCF_001431425.1 | Finkmann et al. 2000 |
| *Stenotrophomonas pavanii* | CBMAI 564; DSM 25135; ICB 89; LMG 25348 | Stems of sugarcane | Brazil | 2011 | GCF_900101175.1 | Ramos et al. 2011 |
| *Stenotrophomonas pictorum* | ATCC 23328; CCM 284; CCUG 1823; CCUG 3368; CIP 103273; DSM 19282; JCM 9942; LMG 981; NCIB 9152; NCIMB 9152; NRRL B-2543; VKM 1240; VKM B-1240 | Soil | unknown | 1928 | GCF_001431585.1 | (Gray and Thornton 1928) Ouattara et al. 2017 |
| *Stenotrophomonas rhizophila* | ATCC BAA-473; CCUG 47042; DSM 14405; e-p10; JCM 13333 | Brassica napus root (rhizosphere oilseed rape) | Germany | 1993 | GCF_000661955.1 | Wolf et al. 2002 |
| *Stenotrophomonas terrae* | DSM 18941; LMG 23958; R-32768 | Soil | Belgium | 2007 | GCF_001431465.1 | Heylen et al. 2007 |
| *Stenotrophomonas tumulicola* | JCM 30961; NCIMB 15009; T5916-2-1b | Viscous gel (biofilm) | Japan | 2016 | GCF_014117215.1 | Handa et al. 2016 |
| *"Stenotrophomonas cyclobalanopsidis"* | CFCC 15341; LMG 31208; TPQG1-4 | Quercus leaves | China | 2018 | GCF_008710035.1 | Bian et al. 2020 |
| *"Stenotrophomonas muris"* | DSM 28631; pT2-440Y | Mouse gut; caecal content; TNFdeltaARE/+ C57BL/6 mouse | Germany | 2009 | GCF_024621935.1 | Afrizal et al. 2022 |

| | | | | | | |
|---|---|---|---|---|---|---|
| *"Stenotrophomonas nematodicola"* | CGMCC19401; CPCC 101271; KCTC XXX; W5 | Soil | China | 2019 | GCF_009467805.1 | Wei et al. 2021 |
| *"Stenotrophomonas panacihumi"* | JCM 16536; KCTC 22893; KEMB 9004-002; MK06 | Soil of a ginseng field | South Korea | 2010 | GCF_001431645.1 | Yi et al. 2010 |
| *"Stenotrophomonas pennii"* | Sa5BUN4 | *Gallus gallus* | UK | 2020 | GCF_014836545.1 | Gilroy et al. 2021 |
| *"Stenotrophomonas sepilia"* | JCM 32102; KCTC 62052; SM16975 | Homo sapiens blood | India | 2012 | GCF_003244875.1 | Gautam et al. 2021 |
| ***Xanthomonas* genus** | | | | | | |
| *Xanthomonas hawaiiensis* sp. nov. | A6251; D-93 | *Spathiphyllum* | USA: Hawaii | 1985 | CP115873 | This study |
| *Xanthomonas hawaiiensis* sp. nov. | *A2111; D-194 | *Colocasia* | USA: Hawaii | 1986 | JAQMHB000000000 | This study |
| *Xanthomonas albilineans* | ATCC 33915; CFBP 2523; DSM 3583; ICMP 196; LMG 494; NCPPB 2969 | *Saccharum officinarum* | Fiji | 1961 | GCF_002939705.1 | (Ashby 1929) Dowson 1943 (Approved Lists 1980) |
| *Xanthomonas arboricola* | ATCC 49083; CFBP 2528; DSM 18808; ICMP 35; LMG 747; NCPPB 411; pv. Juglandis | *Juglans regia* | New Zealand | 1956 | GCF_001013475.1 | Vauterin et al. 1995 |
| *Xanthomonas axonopodis* | ATCC 19312; DSM 3585; ICMP 50; LMG 538; LMG 982; NCPPB 457 | *Axonopus scoparius* | Colombia | 1949 | GCF_001304695.1 | Starr and Garces 1950 (Approved Lists 1980) |
| *Xanthomonas bonasiae* | CFBP 8703; DSM 112530; FX4 | *Ficus benjamina* (Crown gall) | Iran | 2019 | GCA_017163705.1 | Mafakheri et al. 2022 |
| *Xanthomonas bromi* | CFBP 1976; DSM 18804; ICMP 12545; LMG 947 | *Bromus carinatus* | France | 2013 | GCA_900092025.1 | Vauterin et al. 1995 |
| *Xanthomonas campestris* | ATCC 33913; CFBP 2350; CIP 100069; DSM 3586; ICMP 13; LMG 568; NCPPB 528 | *Brassica oleracea* | UK | - | | (Pammel 1895) Dowson 1939 (Approved Lists 1980) |

| | | | | | | |
|---|---|---|---|---|---|---|
| *Xanthomonas cassavae* | DSM 18958; ICMP 204; LMG 673; NCPPB 101 | *Manihot esculenta* | Malawi | 1951 | GCA_000454545.1 | (ex Wiehe and Dowson 1953) Vauterin et al. 1995 |
| *Xanthomonas cissicola* | ATCC 33616; CCUG 18839; CFBP 2432; CIP 106723; DSM 21306; JCM 13362; NCPPB 2982 | *Causonis japonica* | Japan | 1974 | GCF_008801575.1 | (Takimoto 1939) Rudra and Gupta 2021 |
| *Xanthomonas citri* | ATCC 49118; Gabriel 3213; ICMP 15804; ICPB 10518; LMG 9322 | *Citrus aurantiifolia* | USA | 1915 | GCF_002018575.1 | (ex Hasse 1915) Gabriel et al. 1989 |
| *Xanthomonas codiaei* | ATCC 700187; DSM 18812; ICMP 9513; LMG 8678 | *Codiaeum variegatum var. Pictum cv. Superstar* | USA | 1987 | GCA_002939785.1 | Vauterin et al. 1995 |
| *Xanthomonas cucurbitae* | CFBP 2542; DSM 18957; ICMP 2299; LMG 690; NCPPB 2597 | *Cucurbita maxima* | New Zealand | 1968 | GCA_002939885.1 | (ex Bryan 1926) Vauterin et al. 1995 |
| *Xanthomonas dyei* | CFBP 7245; ICMP 12167; NCPPB 4446 | *Metrosideros excelsa* | New Zealand | 1993 | GCA_002939865.1 | Young et al. 2010 |
| *Xanthomonas euroxanthea* | CCOS 1891; CPBF 424; LMG 31037; NCPPB 4675 | *Juglans regia* | Portugal | 2016 | GCA_900476395.1 | Martins et al. 2020 |
| *Xanthomonas euvesicatoria* | ATCC 11633; DSM 19128; ICMP 109; ICMP 98; LMG 27970; NCPPB 2968 | *Capsicum frutescens* | USA | 2007 | GCF_001401555.1 | Jones et al. 2006 |
| *Xanthomonas floridensis* | ATCC TSD-60; ICMP 21312; LMG 29665; NCPPB 4601; WHRI 8848 | *Nasturtium officinale* | USA | 1994 | GCA_001642575.1 | Xanthomonas floridensis Vicente et al. 2017 |
| *Xanthomonas fragariae* | ATCC 33239; CCUG 23372; CFBP 2157; DSM 3587; ICMP 5715; LMG 708; NCPPB 1469; VKM B-2165 | *Fragaria chiloensis var. ananassa* | USA | 1985 | GCA_900183975.1 | Kennedy and King 1962 (Approved Lists 1980) |

| | | | | | | |
|---|---|---|---|---|---|---|
| *Xanthomonas hortorum* | CFBP 5858; DSM 19143; ICMP 453; LMG 733; NCPPB 939; pv. Hederae | *Hedera helix* | USA | 1961 | GCF_003064105.1 | Vauterin et al. 1995 |
| *Xanthomonas hyacinthi* | ATCC 19314; CFBP 1156; DSM 19077; ICMP 189; LMG 739; NCPPB 599 | *Hyacinthus orientalis* | Netherlands | 1958 | GCF_009769165.1 | (ex Wakker 1883) Vauterin et al. 1995 |
| *Xanthomonas hydrangeae* | CCOS 1956; GBBC 2123; LMG 31884 | *Hydrangea arborescens* | Belgium | 2011 | GCA_905142475.1 | Dia et al. 2021 |
| *Xanthomonas maliensis* | CFBP 7942; LMG 27592; M97 | *Oryza sativa* | Mali | 2009 | GCA_009192945.1 | Triplett et al. 2015 |
| *Xanthomonas melonis* | DSM 18798; ICMP 8682; LMG 8670; NCPPB 3434 | *Cucumis melo* | Brazil | 1974 | GCA_002940015.1 | Vauterin et al. 1995 |
| *Xanthomonas nasturtii* | ATCC TSD-61; ICMP 21313; LMG 29666; NCPPB 4600; WHRI 8853 | *Nasturtium officinale* | USA | 2014 | GCA_001660815.1 | Vicente et al. 2017 |
| *Xanthomonas oryzae* | ATCC 35933; CFBP 2532; Dye YK9; ICMP 3125; LMG 5047; NCCPPB 3002; PDDCC 3125; Rao X08 | *Oryza sativa* | Belgium | 1965 | GCA_004136375.1 | (ex Ishiyama 1922) Swings et al. 1990 |
| *Xanthomonas phaseoli* | ATCC 49119; G27; LMG 29033 | *Phaseolus vulgaris* | USA | 1989 | GCA_022749655.1 | (ex Smith 1897) Gabriel et al. 1989 |
| *Xanthomonas pisi* | ATCC 35936; DSM 18956; ICMP 570; LMG 847; NCPPB 762 | *Pisum sativum* | Japan | 1997 | GCF_001010415.1 | (ex Goto and Okabe 1958) Vauterin et al. 1995 |
| *Xanthomonas populi* | ATCC 51165; CFBP 1817; DSM 18847; ICMP 5816; LMG 5743; NCPPB 2959 | *Populus canadensis* | France | 1957 | GCA_002940065.1 | (ex Ridé 1958) Ridé and Ridé 1992 |
| *Xanthomonas prunicola* | CECT 9404; CFBP 8353; IVIA 3287.1 | *Prunus persica var. nectarina* | Spain | 2015 | GCA_002846205.1 | López et al. 2018 |
| *Xanthomonas sacchari* | CFBP 4641; DSM 22617; ICMP 16916; LMG 471 | *Saccharum officinarum* | Guadeloupe | 1980 | GCF_002940085.1 | Vauterin et al. 1995 |

| | | | | | | |
|---|---|---|---|---|---|---|
| *Xanthomonas theicola* | CFBP4691; ATCC 700184; DSM 18797; ICMP 6774; LMG 8684 | *Camellia sinensis* | Japan | 1974 | GCA_014236795.1 | Vauterin et al. 1995 |
| *Xanthomonas translucens* | ATCC 19319; CFBP 2054; DSM 18974; ICMP 5752; LMG 876; NCPPB 973 | *Hordeum vulgare* | USA | 1933 | GCA_900094325.1 | (ex Jones et al. 1917) Vauterin et al. 1995 |
| *Xanthomonas vasicola* | CFBP 2543; DSM 16926; ICMP 3103; LMG 736; NCPPB 2417 | *Sorghum bicolor* | New Zealand | 1969 | GCA_000772705.2 | Vauterin et al. 1995 |
| *Xanthomonas vesicatoria* | ATCC 35937; CFBP 2537; DSM 22252; ICMP 63; LMG 911; NCPPB 422 | *Solanum lycopersicum* | New Zealand | 1955 | GCF_001908725.1 | (ex Doidge 1920) Vauterin et al. 1995 |
| *Xanthomonas youngii* | AmX2T; CFBP 8902; DSM 112529 | *Amaranthus sp. (Crown gall)* | Iran | 2019 | GCA_017163755.1 | Mafakheri et al. 2022 |
| *"Xanthomonas cannabis"* | NCPPB 2877 | *Cannabis sativa* | Romania | 1974 | GCF_000802365.1 | Jacobs et al. 2015 |
| *"Xanthomonas indica"* | CFBP 9039; ICMP 24394; MTCC 13185; PPL560T | Rice seeds | India | 2021 | GCA_022669045.1 | Rana et al. 2022 |
| *"Xanthomonas massiliensis"* | CSUR P2129; SN 8 | Stools of obese patient | France | 2015 | GCF_900018785.1 | Ndongo et al. 2017 |
| *"Xanthomonas sontii"* | CFBP 8688; ICMP 23426; JCM 33631; MTCC 12491; PPL1 | *Oryza sativa* | India | 2012 | GCA_008119715.1 | Bansal et al. 2021 |
| *"Xanthomonas surreyensis"* | Sa3BUA13 | *Gallus gallus* | UK | 2020 | GCF_014836395.1 | Gilroy et al. 2021 |

* indicates the strain is not type strain. The species name between two ditto marks (“) indicates the invalidly published species.

**TABLE S2**. Antibiotic sensitivity assays of new species from Araceae, *Xanthomonas hawaiiensis* sp. nov., *Stenotrophomonas aracearum* sp. nov., and *S. oahuensis* sp. nov., using seven antibiotics.

| Bacteria species | Strains | Bacitracin (50 mg/ml) | Chloramphenicol (50 mg/ml) | Gentamicin (50 mg/ml) | Kanamycin (50 mg/ml) | Penicillin (50 mg/ml) | Tetracycline (40 mg/ml) | Polymyxin B Sulfate (50 mg/ml) |
|---|---|---|---|---|---|---|---|---|
| *Xanthomonas hawaiiensis* sp. nov. | $A6251^T$ | + | +++ | ++ | ++ | +++ | +++ | + |
| *Xanthomonas hawaiiensis* sp. nov. | A2111 | + | +++ | +++ | ++ | +++ | +++ | +++ |
| *Stenotrophomonas aracearum* sp. nov. | $A5588^T$ | + | ++ | ++ | ++ | - | ++ | + |
| *Stenotrophomonas oahuensis* sp. nov. | $A5586^T$ | -/+ | ++ | ++ | ++ | + | ++ | + |

(-: no inhibition zone; -/+: size in radius = 0.1 cm; +:0 < size in radius ⩽ 1.0 cm; ++: 1.0 cm < size in radius ⩽ 2.0 cm; +++: 2.0 cm < size in radius)
